# Neural Heterogeneity Enhances Reliable Neural information Processing: Local Sensitivity and Globally Input-slaved Transient Dynamics

**DOI:** 10.1101/2024.06.30.599443

**Authors:** Shengdun Wu, Haiping Huang, Shengjun Wang, Guozhang Chen, Changsong Zhou, Dongping Yang

## Abstract

Cortical neuronal activity varies over time and across repeated stimulation trials, yet consistently represents stimulus features. The dynamical mechanism underlying this reliable representation and computation remains elusive. This study uncovers a mechanism that achieves reliable neural information processing, leveraging a biologically plausible network model with neural heterogeneity. We first investigate neuronal timescale diversity in reliable computation, revealing it disrupts intrinsic coherent spatiotemporal patterns, enhances local sensitivity, and aligns neural network activity closely with inputs. This leads to local sensitivity and globally input-slaved transient dynamics, essential for reliable neural processing. Other neural heterogeneities, such as non-uniform input connections and spike threshold heterogeneity, plays similar roles, highlighting neural heterogeneity’s role in shaping consistent stimulus representation. This mechanism offers a potentially general framework for understanding neural heterogeneity in reliable computation and informs the design of new reservoir computing models endowed with liquid wave reservoirs for neuromorphic computing.

**Teaser:** Neural diversity disrupts spatiotemporal patterns, aligning network activity with inputs for reliable information processing.

## 1. Introduction

Cortical neurons exhibit substantial spiking time irregularity and trial-to-trial variability in both spontaneous activities and evoked responses to repeated stimuli [1, 2]. Despite this notable variability, neural representations of stimuli remain functionally consistent [3–6]. Stimuli onset widespreadly quenches neural variability and leads to reliable sensory coding [6, 7]. Neural population dynamics has been suggested to straightforwardly produce resilient movement patterns, even when confronted with highly unpredictable neural responses [8, 9]. Thus, a central challenge is to understand the neural mechanisms that reconcile inherent cortical variability with reliable representations of external inputs.

Randomly connected recurrent neural networks (RNNs) with excitation-inhibition (E-I) balance [10] are often used to model irregular and asynchronous activities with reliable macroscopic dynamics, but mainly track network input [8, 10, 11]. Spatially extended spiking neural networks (SNNs), incorporating the distance-dependent connection probability [12], limit the dynamical complexity but can also generate intricate spatiotemporal dynamics through Turing-Hopf bifurcation that produces coherent spatiotemporal patterns [13, 14]. Such destabilized networks can be used for reservoir computing [15], whose reliability can be achieved by breaking the symmetry of network dynamics via non-uniform input connections [13], referred to as neural heterogeneity in terms of heterogeneous input connections. It indicates neural heterogeneity essentially contributes to neural computation, as previously demonstrated, *e*.*g*., through the heterogeneity of cell spiking thresholds [16] or membrane time constants [17]. However, the underlying dynamical mechanism remains elusive.

Biologically plausible RNNs can provide insights into how biological networks perform reliable computational functions, *e*.*g*., encoding, decoding and learning [16]. The involved neural populations are highly heterogeneous, varying in structure, gene expression, and electrophysiological properties [18–20], such as membrane capacitance and resistance, resting potential, or spiking threshold. These properties capture variations in the structural composition of the cell membrane across neurons of the same type as well as distinct types [18–20]. Emerging research suggests that neuronal diversity plays a pivotal role in information processing [16, 17, 21–24]. Remarkably, the comprehensive large-scale model [22] of V1 area assimilating anatomical and neurophysiological data [18] can solve diverse visual processing tasks with higher energy efficiency and greater robustness to noise than current artificial intelligence models. Even within simplified SNNs lacking a detailed anatomical architecture, neuronal heterogeneity in terms of membrane and synaptic time constants (observed in the brain [18–20]) can enhance robust learning [17]. However, they lack a dynamical mechanism to understand or even unify the roles of various neural heterogeneities.

Here, we reveal a potentially general dynamical mechanism, starting from investigating two specific neural diversities: heterogeneous leakage time constants and response time constants [18, 25]. We explore their roles in computation’s reliability regarding an input-output mapping task, employing a biologically plausible SNN model. Our findings show that either diversity can enhance reliable computation, playing a similar role to that of non-uniform input connections [13]. It suggests a unique dynamical mechanism, relying on consistent representation: the diversity disrupts intrinsic coherent spatiotemporal patterns [13, 26], leading to local sensitivity and globally input-slaved transient dynamics [27]. This dynamics shapes the high-dimensional representation, inducing a liquid-like spatiotemporal activity pattern and forming input-slaved trajectories for reliably representing input information [27]. This mechanism is robust across networks with varying connection ranges and connectivity randomness, and explains the similar roles of others: non-uniform input connections or spike threshold heterogeneity. It provides a potentially uniform framework for understanding the significance of neural heterogeneity in ensuring reliable neural information processing. Thus, our work sheds light on new reservoir computing models endowed with this mechanism.

## 2. Results

To investigate the dynamical mechanism underlying reliable computation, we start from exploring the effect of neuronal timescale diversity on spatiotemporal activity properties in Sec. 2.1, leveraging the spatially extended E-I SNNs with current-based leaky integrate-and-fire (LIF) neurons (Fig. 1(A)). Specifically, we first consider the diversity of two time constants in neuronal dynamics: leakage time constant *τ*_*L*_ and response time constant *τ*_Γ_ (Fig. 1(B)). Section 2.2 shows that neuronal timescale diversity can lead to reliable computation and representation with no need for fine-tuning connection ranges in Sec. 2.2. In Sec. 2.3, the dynamical mechanism is presented and comprehended from three perspectives: local responsive sensitivity, input-induced global spatiotemporal modes and high-dimensional transient dynamics, ensuring robust and reliable representation and computation. In Sec. 2.4, this mechanism also explains the similar roles of other neural heterogeneities, such as non-uniform input connections and spike threshold heterogeneity, demonstrating a potentially general framework for understanding the role of neural heterogeneity in ensuring reliable neural information processing.

**Fig. 1.**
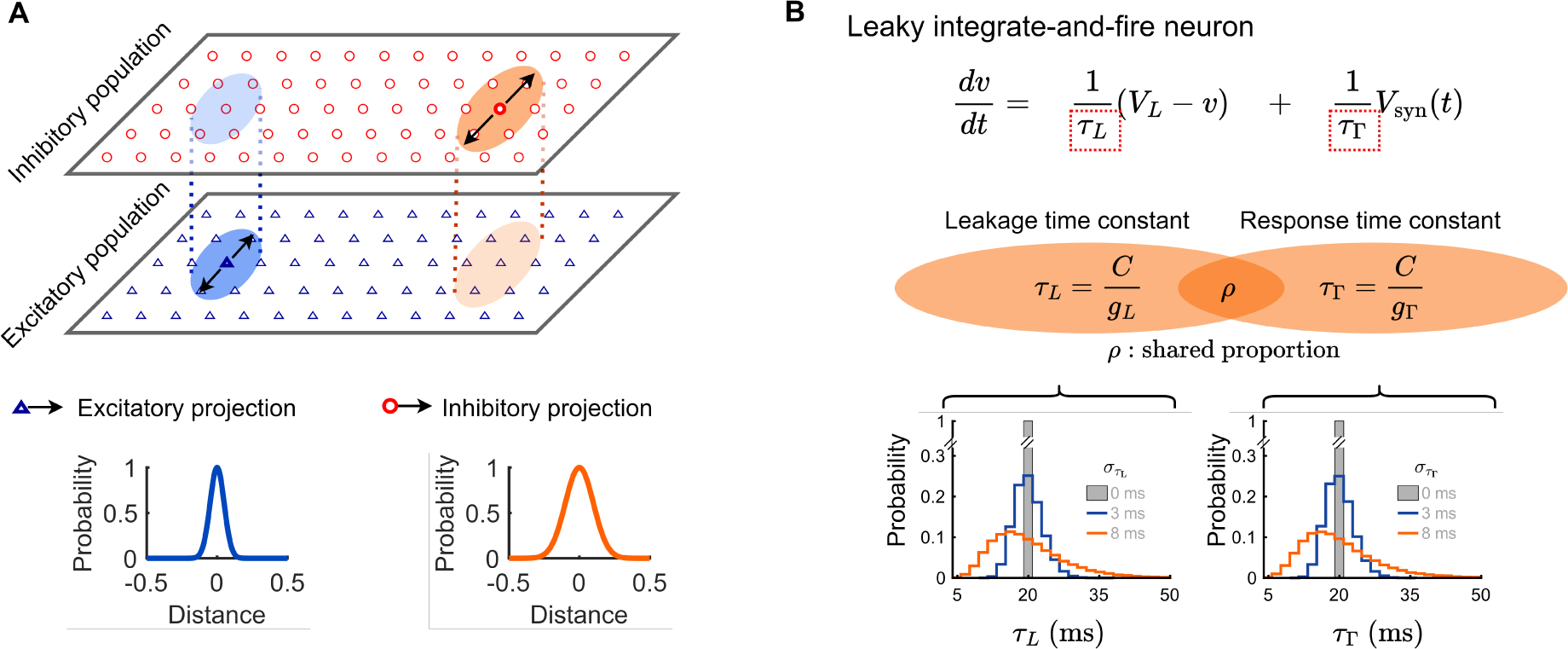
Schematic of spatially extended SNNs with diversity of two time constants in neuronal dynamics. (**A**) Network architecture. Excitatory (E, orange) and inhibitory (I, blue) neurons evenly distribute on a two-dimensional plane: [0, 1] × [0, 1]. They are connected with a distance-dependent Gaussian probability, characterized by connection ranges *σ*_*e*_ and *σ*_*i*_ for E and I neurons, respectively. By default, *σ*_*e*_ = 0.05 and *σ*_*i*_ = 0.1. (**B**) LIF neuron model and time constants’ distributions. Top: Illustration of the current-based LIF neuron model, where *τ*_*L*_ and *τ*_Γ_ are leakage and response time constants, and *V*_syn_ represents the overall postsynaptic potential. Middle: Definition of time constants *τ*_*L*_ and *τ*_Γ_ with their correlation *ρ*. Bottom: Distributions of time constants *τ*_*L*_ and *τ*_Γ_ with widths 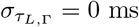 (grey), 3 ms (blue), and 8 ms (orange).

### 2.1. Model Setting and Effect of Neuronal Timescale Diversity on Spatiotemporal Dynamics

We commence by illustrating in Fig. 1(A) the spatially extended E-I SNN with distance-dependent connection probabilities [13, 28]. The connectivity profiles can be characterized by connection ranges *σ*_*e*_ and *σ*_*i*_ for E and I neurons, respectively. Broader recurrent inhibition (*σ*_*i*_ *>* 2*σ*_*e*_) can lead to pattern formation through a Turing-Hopf bifurcation: breaking the spatiotemporal symmetry and the local E-I balance, as demonstrated by mean-field analysis under a diffusion approximation [28] with the ansatz of Gaussian distribution of firing rates [13] (Fig. 2(D), blue). It results in a remarkably coherent spatiotemporal pattern of collective activities of E neurons, wherein local E populations exhibit synchronized oscillations and spiking activity is irregular (Fig. 2(A and B)). It spatially destabilizes the local E-I balance, leading to pronounced temporal fluctuations in firing rates (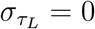 in Fig. 2(C)).

**Fig. 2.**
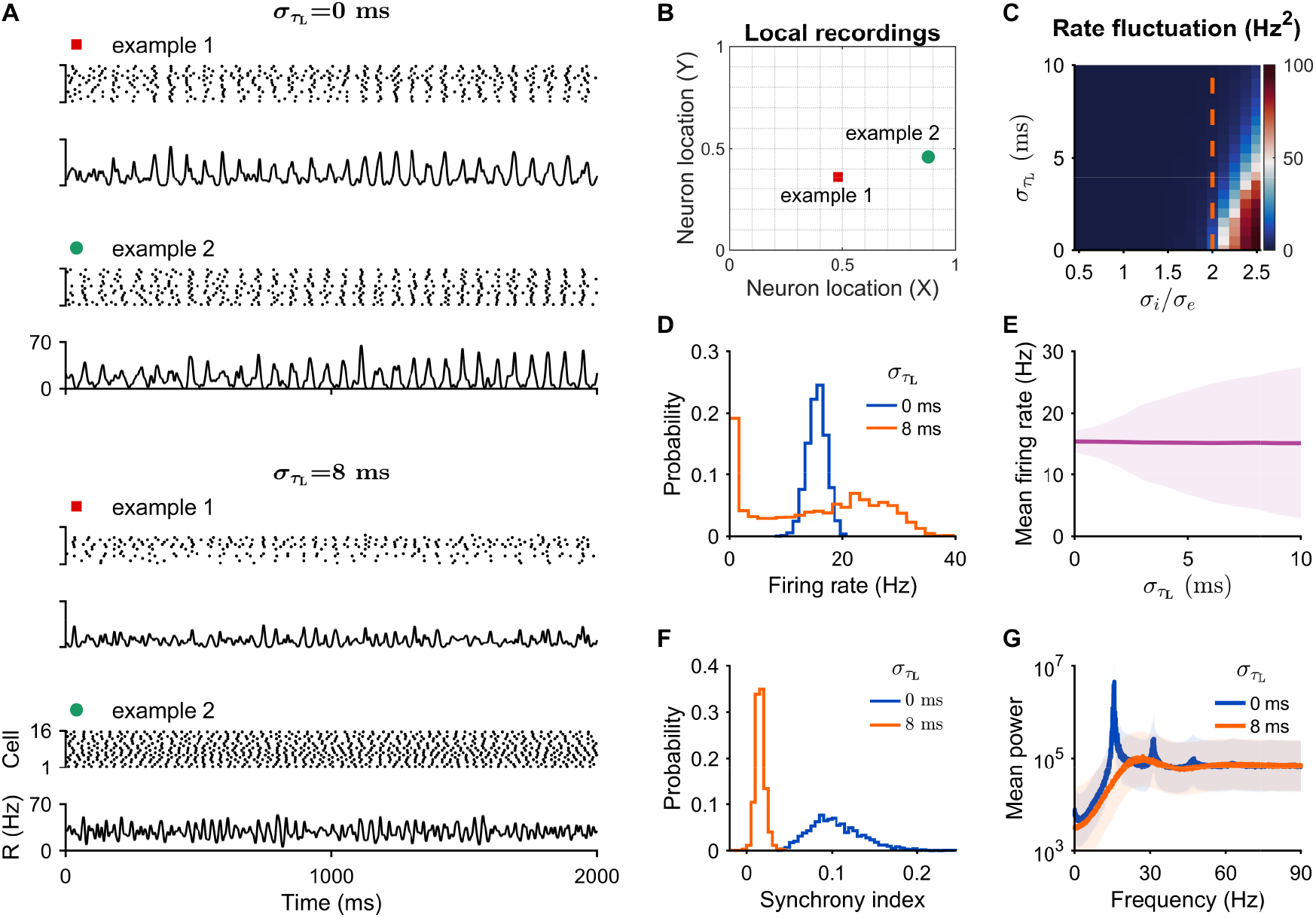
Spontaneous spatiotemporal dynamics with/without timescale diversity. We record activity of 2500 spatial sites with each one comprising 16 E neurons. (**A**) Raster plot and firing rate series of two randomly chosen sites in the homogeneous network (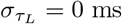, top) and the heterogeneous network (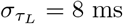, bottom). (**B**) Spatial positions of the two sites. (**C**) Temporal fluctuation of E neurons’ firing rates across the parameter space 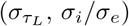. (**D**) Firing rate distributions of E neurons at 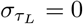 ms or 8 ms. (**E**) Mean firing rates (thick line) and their standard deviations (SDs, shaded area) plotted against 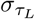. (**F**) Synchrony index distributions. (**G**) Average power spectra (thick lines) of individuals’ firing rates at 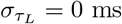 or 8 ms, with shaded areas denoting SDs.

Under this network, we consider two distinct types of neuronal time constants in the LIF neuron model (Fig. 1(B)): The leakage time constant *τ*_*L*_ controls the leak rate of the neuron’s membrane potential; The response time constant *τ*_Γ_ regulates the overall scale of network drives [18, 25]. Here, we consider scenarios where *τ*_*L*_ and *τ*_Γ_ exhibit diversity and correlation, as evidenced in the biological datasets [18, 25]. As the timescale is non-negative, the diversity is introduced with simple lognormal distributions, akin to those measured in experimental data [19, 20], and characterized by distribution widths 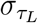 and 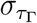 for *τ*_*L*_ and *τ*_Γ_ (Fig. 1(B)), respectively. The parameter *ρ* controls the shared proportion between the distributions of *τ*_*L*_ and *τ*_Γ_. When *ρ* = 1, *τ*_*L*_ and *τ*_Γ_ are completely correlated, both equal to the membrane time constant *τ*_*m*_, and the neuronal model is equivalent to that used in [17]. For more details, refer to Sec. 5.

Now we investigate the effects of *τ*_*L*_ diversity on network dynamics (Fig. 2). Increasing *τ*_*L*_ diversity gradually disrupts Turing-Hopf bifurcation [13, 26], decreasing temporal rate fluctuations (Fig. 2(C)) and leading to more robust asynchronous states [11]. With *τ*_*L*_ diversity, neuronal firing rates exhibit a broader non-Gaussian distribution of firing rates (Fig. 2(D), orange): a weighted summation of timescale-dependent Gaussian distributions (fig. S1), thus complicating the mean-field analysis of this bifurcation. Nonetheless, it can be understood via a simplified model of diffusively coupled nonlinear oscillators, where timescale diversity has been proved to stabilize the asynchronous state and prevent interaction-induced coherence [29–31]. Specifically, a larger 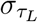 induces a larger heterogeneity of neuronal firing rates, while its overall average remains stable (Fig. 2(E)). In the heterogeneous network 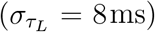, neuronal firing is irregular, and different recording sites show different activity levels (Fig. 2(A)). Such spatial heterogeneity in neuronal activity is consistent with experimental records showing considerable variability in firing rates across neurons [32]. Compared to the homogeneous network 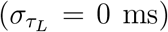, neurons in this heterogeneous network exhibit various weaker oscillatory patterns (Fig. 2(A)), leading to significantly lower pairwise synchrony (Fig. 2(F)) and much weaker average oscillatory amplitudes with a broader frequency distribution (Fig. 2(G)).

Similar effects of *τ*_Γ_ diversity on network dynamics are presented in fig. S2. With either timescale diversity, neurons’ linear response functions behave spatially heterogeneous, and thus their diversities can disrupt intrinsic coherent spatiotemporal patterns. Such destabilized network dynamics have been expected to support reliable computation [13].

### 2.2. Timescale Diversity Enhancing Reliability of Input-output Mapping Computation and Mackey-Glass Signal Representation

We now examine the role of timescale diversity in reliable computation regarding input-output mapping in a reservoir computing setting (Fig. 3(A)). In this task, the network is driven by a sinusoidal input and local rate readouts as the “reservoir” is linearly combined to produce output time series (Fig. 3(A and B)). Readout weights are trained using a ridge least-square method (LSM) to mold the output to target time series (Fig. 3(A and C)). In the homogeneous network (*τ*_*L*,Γ_ = 0), the outputs poorly track the target time series and display noticeable variability across trials (Fig. 3(C)). In contrast, both *τ*_*L*_ and *τ*_Γ_ diversity can reliably produce outputs closely matching the target (Fig. 3(C)). These two diversities consistently enhance the computation’s reliability, as larger 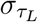 or 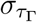 leads to lower normalized root-mean-square error (NRMSE) across multiple trials (Fig. 3(D and E)), indicating a reduced trial-to-trial variability.

**Fig. 3.**
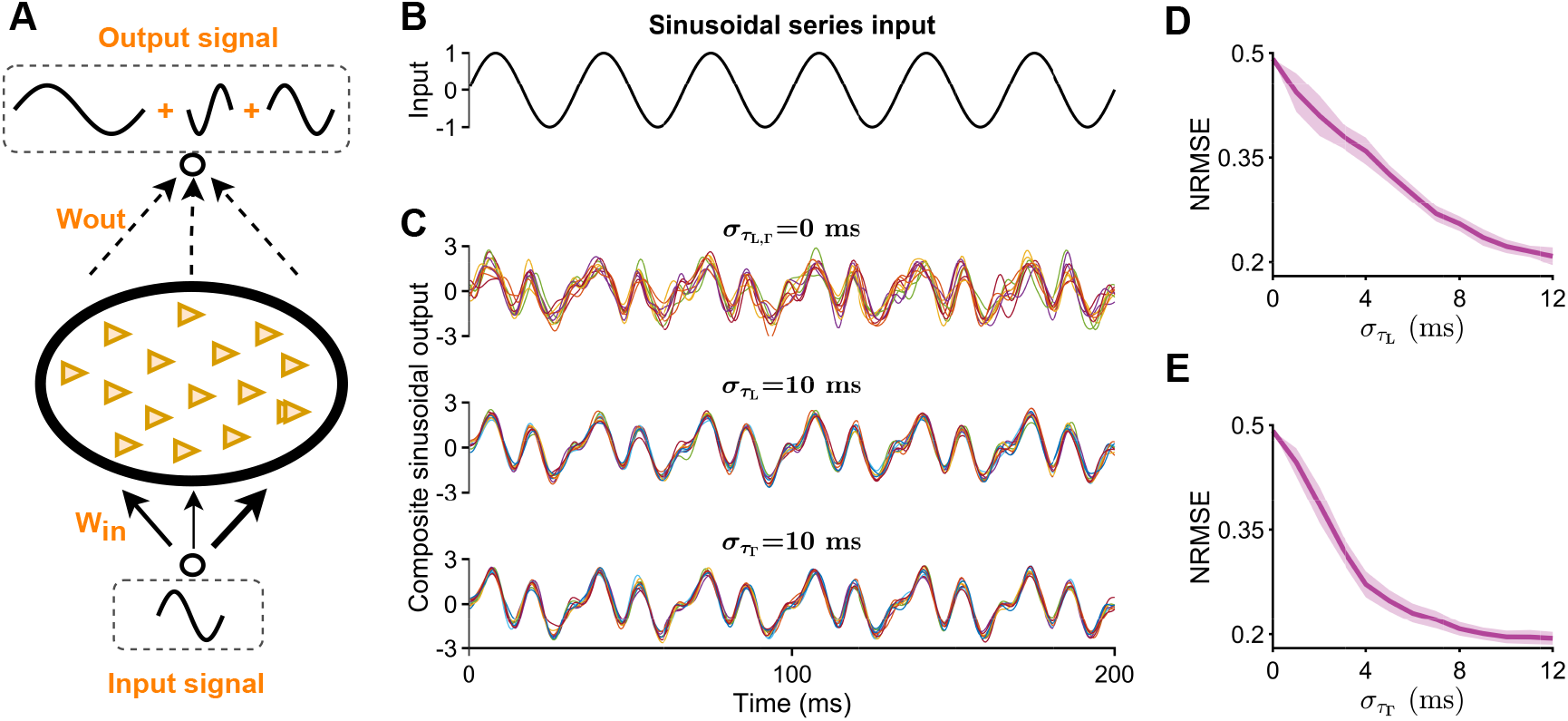
Enhanced reliability of computation. (**A**) Schematic of the input-output mapping task. The input signal is sinusoidal: *I*_sin_ = *ϵ* · sin(2*πf*_sin_*t*), and the target output is a composite of three sinusoidal signals: *I*_target_ = 1.5 · [sin(*πf*_sin_*t*) + sin(2*πf*_sin_*t*) − sin(3*πf*_sin_*t*)]. (**B**) Time trace of the input. (**C**) Computations from: the homogeneous network (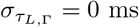, top), the heterogeneous networks with leakage timescale diversity (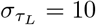 ms, middle), or response timescale diversity (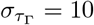 ms, bottom). (**D** and **E**) Target fitting normalized root-mean-square error (NRMSE) plotted against 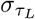 (D) and 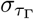 (E). In panel (C), the black line represents the target signal, and the overlapped colored lines depict the outputs from 10 repeated trials. Parameter setting: amplitude *ϵ* = 30 mV and frequency *f*_sin_ = 30 Hz.

Such reliable computation in heterogeneous networks reminds us of a reliable information representation. To examine this, we repeatedly apply a segment of one-dimensional Mackey-Glass (MG) chaotic series [33] to the networks and utilize the LSM method to reconstruct the input from neural response activities (Fig. 4(A)). MG signal is characterized by its intrinsic pace *f*_MG_ [33], which showcases a distinct frequency range (Fig. 4(B)). As depicted in Fig. 4(A), the reconstructed MG output signals from 10 repeated trials (overlapping colored lines) closely follow the trend of the MG input signal (black line) but exhibit various degrees of variability across trials. For the homogeneous network 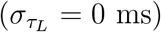, the outputs show considerable trial-to-trial variability and are short of accuracy in tracking input signals (Fig. 4(B)). Conversely, for heterogeneous networks with intermediate and high degrees of timescale diversity (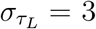 ms and 8 ms), output signals display significantly reduced trial-to-trial variability (Fig. 4(B)). Various MG series with different paces: *f*_MG_ = 20, 50 and 110 can be reliably represented in the heterogeneous network activity (Fig. 4(C)), with a slightly higher NRMSE for the fast-changing MG signal (*f*_MG_ = 110).

**Fig. 4.**
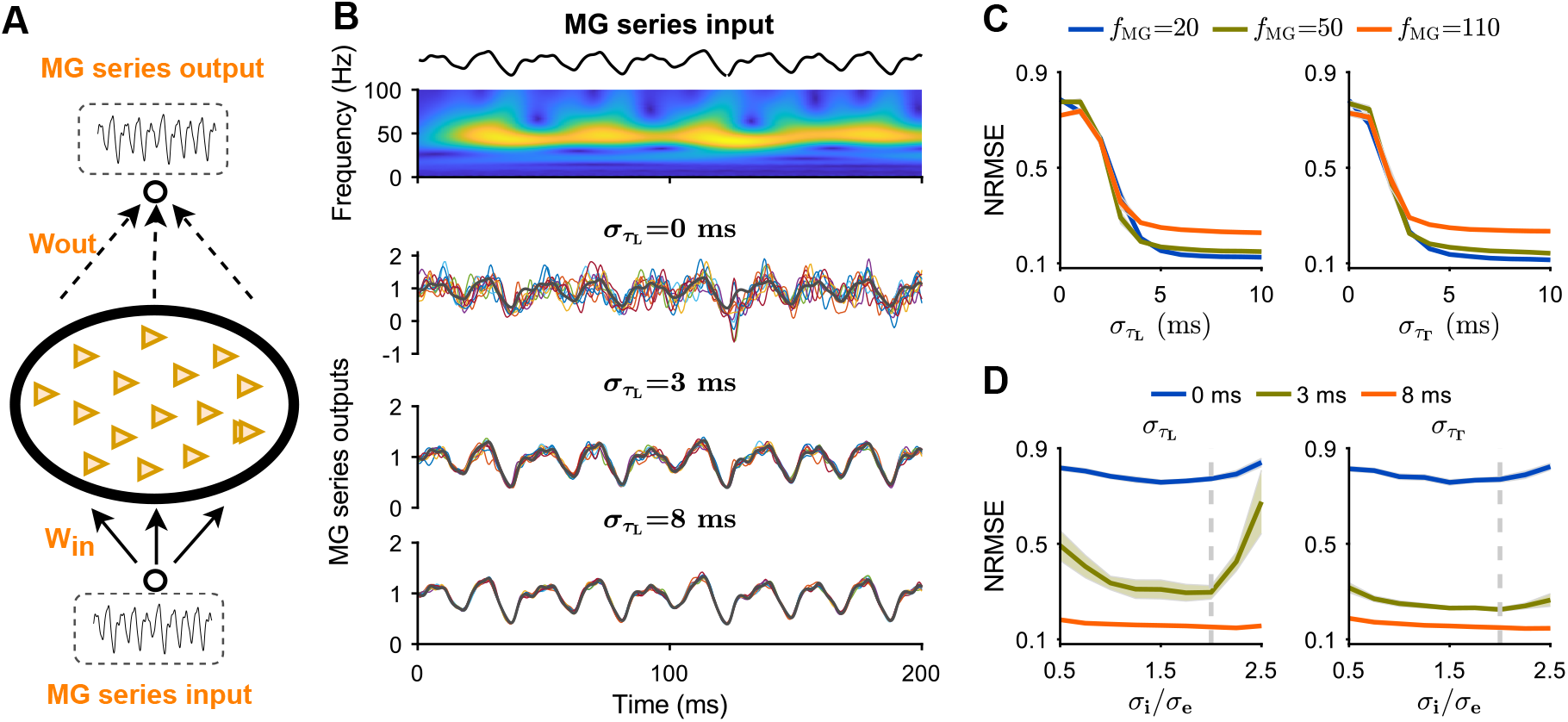
Impact of timescale diversity on the reliability of MG signal reconstruction. (**A**) Task illustration. (**B**) Top: MG signal and its corresponding time-frequency spectrum; Bottom: Reconstructed MG series from networks with 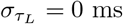, 3 ms, or 8 ms. Line colors denote the same as those in Fig. 3(C). (**C**) NRMSE plotted against 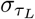 (left) and 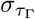 (right) at *f*_MG_ = 20, 50, or 110. (**D**) NRMSE plotted against the ratio *σ*_*i*_*/σ*_*e*_ at 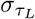(or 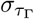) = 0 ms, 3 ms, and 8 ms. Shaded areas represent SDs. In panels (B) and (D), *f*_MG_ = 50.

The representation reliability is robust against various connection range ratios *σ*_*i*_*/σ*_*e*_ (Fig. 4(D)). For homogeneous networks 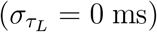 or networks with a low level of timescale diversity 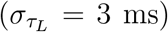, NRMSE initially decreases with increasing *σ*_*i*_*/σ*_*e*_, reaching a minimum before rising again (Fig. 4(D), left). It is due to stabilized rich activity patterns near the critical point (*σ*_*i*_*/σ*_*e*_ ≈ 2) [13]. Conversely, for heterogeneous networks with 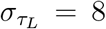 ms, NRMSE remains stable at considerably low values across a wide range of *σ*_*i*_*/σ*_*e*_, demonstrating remarkable robustness against *σ*_*i*_*/σ*_*e*_, as network dynamics remain stable across a broad range of *σ*_*i*_*/σ*_*e*_ (Fig. 2(C)). Even in networks with connections randomly shuffled, both timescale diversity consistently enhance reliable representation (fig. S3).

Similar results are obtained for 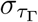 (Fig. 4(C and D), right), demonstrating that timescale diversity essentially enhances reliable representation and computation. It underscores the significant role of timescale diversity, beyond the claimed role of spatially coherent dynamics in overcoming the difficulty of reservoir computing with spiking networks [13]. Its robustness informs a unique underlying dynamical mechanism supporting reliability and robustness.

### 2.3. Dynamical Mechanism of Reliable Representation and Computation

The dynamical mechanism discovered here can be unfolded as the following three perspectives: local responsive sensitivity (Sec. 2.3.1), input-induced global spatiotemporal modes (Sec. 2.3.2) and high-dimensional transient trajectories (Sec. 2.3.3).

#### 2.3.1. Timescale diversity enhancing response sensitivity

We first show how timescale diversity affects the network’s response to external stimuli. We apply small-amplitude sinusoidal periodic signals to the networks and examine the responsive properties (Fig. 5(A)). Both homogeneous and heterogeneous networks exhibit a response power peak at the stimulus frequency *f*_sin_ = 10 Hz or 30 Hz (Fig. 5(B)). The latter shows a higher response amplitude, indicating superior sensitivity to periodic stimuli.

**Fig. 5.**
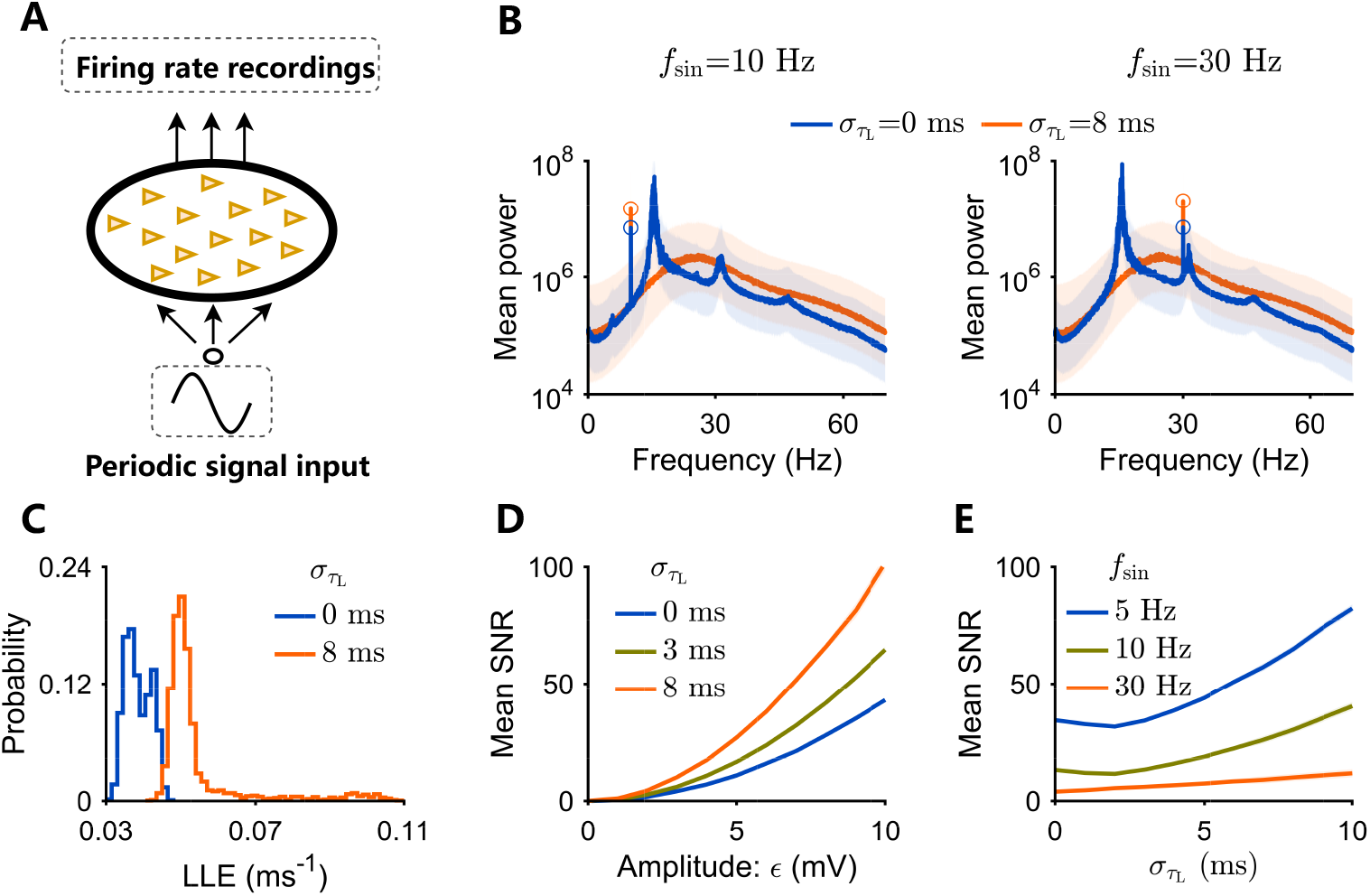
Timescale diversity enhances network responsive sensitivity. (**A**) Schematic representation of the network receiving a sinusoidal input. (**B**) Averaged power spectrum of evoked firing rates for homogeneous (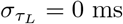, blue) and heterogeneous (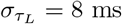, orange) networks. Circular markers denote the peak power spectral density of network’s response to sinusoidal periodic stimuli with an amplitude *ϵ* = 3 mV at *f*_sin_ = 10 Hz (left) or 30 Hz (right). (**C**) Distributions of the largest Lyapunov Exponent (LLE) for homogeneous and heterogeneous networks, calculated from site firing rates without external stimuli. (**D**) Mean Signal-to-Noise Ratio (SNR) plotted against stimulus amplitude *ϵ* at 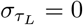 ms, 3 ms, or 8 ms. (**E**) Mean SNR plotted against 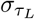 for sinusoidal stimuli at *f*_sin_ = 5 Hz, 10 Hz, or 30 Hz. In panels (B), (D), and (E), shaded areas are SDs.

The superior sensitivity can be attributed to two factors arising from timescale diversity. First, as studied in Sec. 2.1, it can disrupt the intrinsic coherent spatiotemporal patterns [13, 26] (Fig. 2 (A to C) and fig. S4), rendering the site activity more flexible with much higher Largest Lyapunov Exponents (LLE) (Fig. 5(C)). Second, due to the disruption, the local E-I balance recovers (fig. S5), leading to increased sensitivity. This is also evident from the average signal-to-noise ratio (SNR, see Sec. 5), which significantly increases with 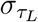 (Fig. 5(D and E)). Thus, timescale diversity improves the effectiveness of network’s response to external stimuli.

Besides, the sensitivity property explains why the correlation between these two timescales impedes the beneficial effects of timescale diversity on computation’s reliability (fig. S6). The leaky timescale *τ*_*L*_, when larger, implies slower leakage and easier firing, leading to a higher firing rate (fig. S6(A), top-left). Conversely, a higher value of *τ*_Γ_ indicates that the neuron takes a longer time to assimilate the driving current, making it hard to fire and resulting in a lower firing rate (fig. S6(A), bottom-left). For both cases, the most sensitive neurons are those firing at a moderate rate (fig. S6(A), right) and thus mismatch in the timescale ranges (fig. S6(B and C)). Consequently, increasing the correlation *ρ*(*τ*_*L*_, *τ*_Γ_) reduces the sensitivity of these neurons (fig. S6(D)). As expected, this decrease in sensitivity diminishes the network’s computation performance (fig. S6(E)).

These results show that while timescale diversity introduces spatial heterogeneity in network activity, it enhances the network’s ability to respond to external signals. It indicates that the dynamics in networks with timescale diversity are more prone to contain the input information. Thus, it can benefit representation and computation.

#### 2.3.2. Timescale diversity shaping consistent stimulus representation

Next, we present how networks represent the external input and how timescale diversity shapes the stimulus representation (Fig. 6(A)). When the input drives networks, the intrinsic spatiotemporal dynamics are perturbed and there is a competition between the intrinsic dynamical modes and the input-induced ones. To demonstrate such competition, we examine both spontaneous and input-induced spatiotemporal activity patterns (Fig. 6(B and D)). The spatiotemporal activity can be viewed as a fluid, and thus the data-driven Dynamic Mode Decomposition (DMD) method [34] can be employed to decompose the high-dimensional neural activities into spatiotemporal dynamical modes (Fig. 6(A)). The dominant dynamical modes should express the input information for reliable representation and computation.

**Fig. 6.**
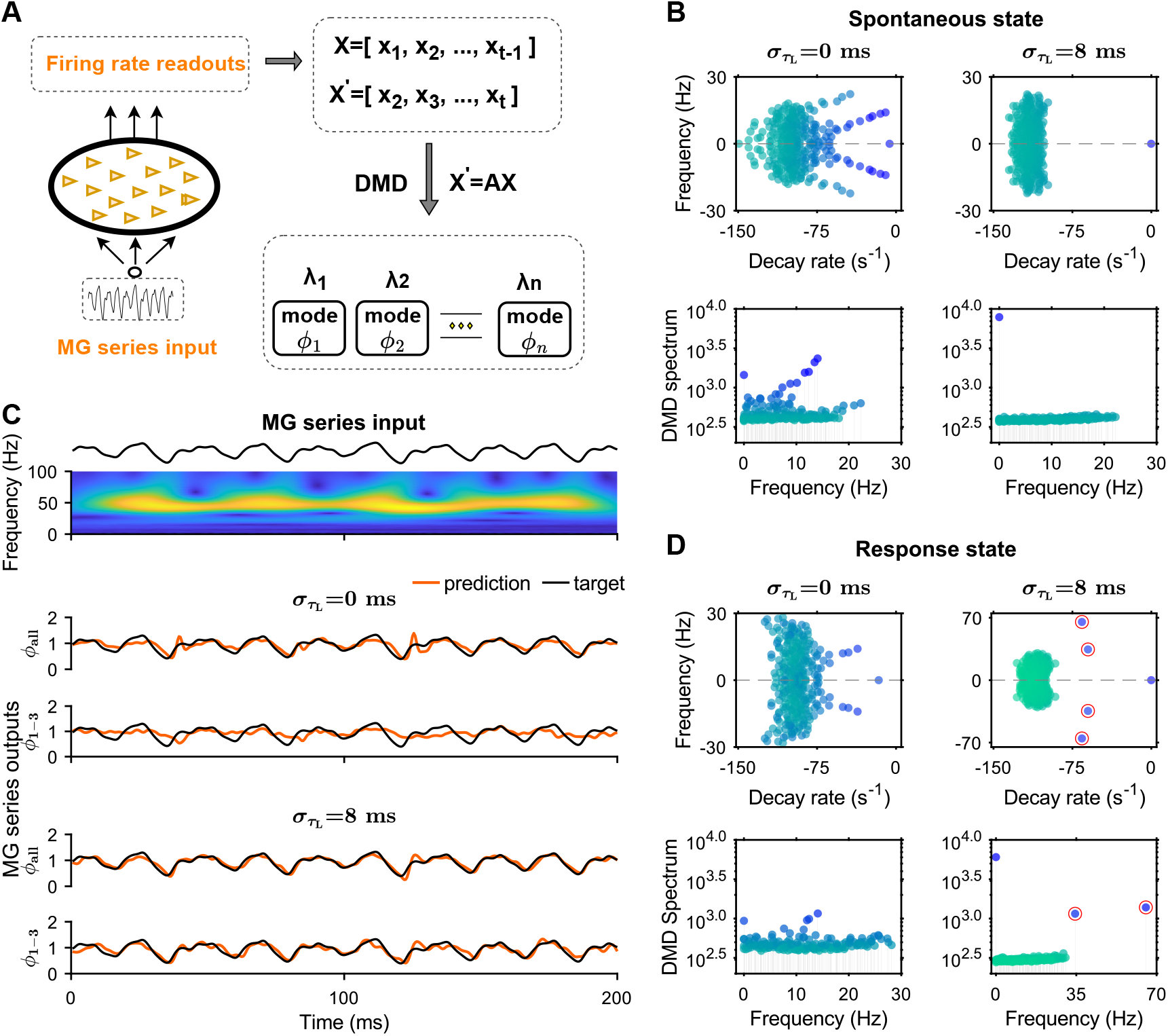
Dynamical mode decomposition (DMD) for characterizing spatiotemporal patterns of neural activities. (**A**) Schematic of the network receiving MG series input and DMD analysis procedure. (**B**) DMD eigenvalue spectrum for spontaneous network activities in homogeneous (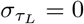 ms, left) or heterogeneous (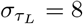, right) networks. Top: DMD eigenvalues *λ* in the complex plane of the decay rate *σ* = Re(ln *λ*_*k*_) · *dt*^−1^ and the frequency *f* = Im(ln *λ*_*k*_) · (2*π* · *dt*)^−1^ with time resolution *dt* = 1 ms.; Bottom: Mode amplitudes (dot colors) plotted against frequencies. (**C**) DMD mode extraction for prediction. Top: MG input signal and corresponding time-frequency spectrum. Bottom: Predicted MG output signals from all modes *ϕ*_all_ or three dominant DMD modes *ϕ*_1-3_. (**D**) Similar to (B), but for network activities in response to MG series input with two new input-induced modes (red circled blue dots).

We demonstrate the role of timescale diversity in shaping consistent stimulus representation by examining the eigenvalue spectra of DMD modes (Sec. 5). As consistent with the Turing-Hopf bifurcation in the homogeneous network (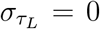 ms), its spontaneous activities show strong intrinsic dynamical modes (Fig. 6(B), left). Applying MG series to the network suppresses its intrinsic modes without inducing a new mode (Fig. 6(D), left). This indicates that the homogeneous network can not generate spatiotemporal patterns matching the frequency range of the MG signal and thus its neural activity lacks the input information. However, in the heterogeneous network 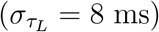, timescale diversity disrupts the intrinsic dynamical modes [13, 26], leaving only one stationary pattern (Fig. 6(B), right), which is the network-wide baseline response to the external input. Interestingly, the input drives this network to induce two new dynamical modes (Fig. 6(D) right, red circled blue dots), which are positioned within the MG frequency range and thus may be enough to encode the input.

We further check whether the induced dynamical modes are enough to represent the external input in the heterogeneous network (Fig. 6(C)). We utilize all modes or three dominant ones to predict the system’s short-term future states (4 time steps ahead). The output weights are linearly regressed to reconstruct the MG input signal from the predicted states. The result confirms that the three dominant modes contain enough information of the input to fulfil this task, whose performance is comparable with that using all modes (Fig. 6(C)). This indicates that these three modes capture the key features of the input, underscoring that the input can be represented in a three-dimensional space. In contrast, in the homogeneous network, even with all modes, precisely reconstructing the MG signal is unavailable, indicating that the network fails to capture the key features of the input (Fig. 6(C)).

These results show that since the heterogeneous network is far from the Turing-Hopf bifurcation and lacks intrinsic coherence (Fig. 2(C)), it effectively responds to external stimuli and captures the features of stimulus signals. It demonstrates that timescale diversity shapes consistent stimulus representation by inducing input-related dynamical modes, thanks to its locally enhanced sensitivity to external inputs. Therefore, it is beneficial for reliable representation and computation.

#### 2.3.3. Timescale diversity reducing high-dimensional trial-to-trial variability

Finally, we show that the input can be reliably and robustly represented by the heterogeneous network activity in a three-dimensional space. For both homogeneous and heterogeneous networks, local rates approximately track the shared input with the addition of irregular fluctuations, consistent with Poisson-like spike-timing variability (Fig. 4(A)). Applying principal component (PC) analysis to the local rates, we extract the intrinsic low-dimensional structure of population response activities (Fig. 7(A)). It reveals that the majority of firing rate variability is captured by the first PC projection, representing the variability inherited from the external input MG series (Fig. 7(A and B)). The remaining variability spreads among higher PC projections representing spatially unstructured variability.

**Fig. 7.**
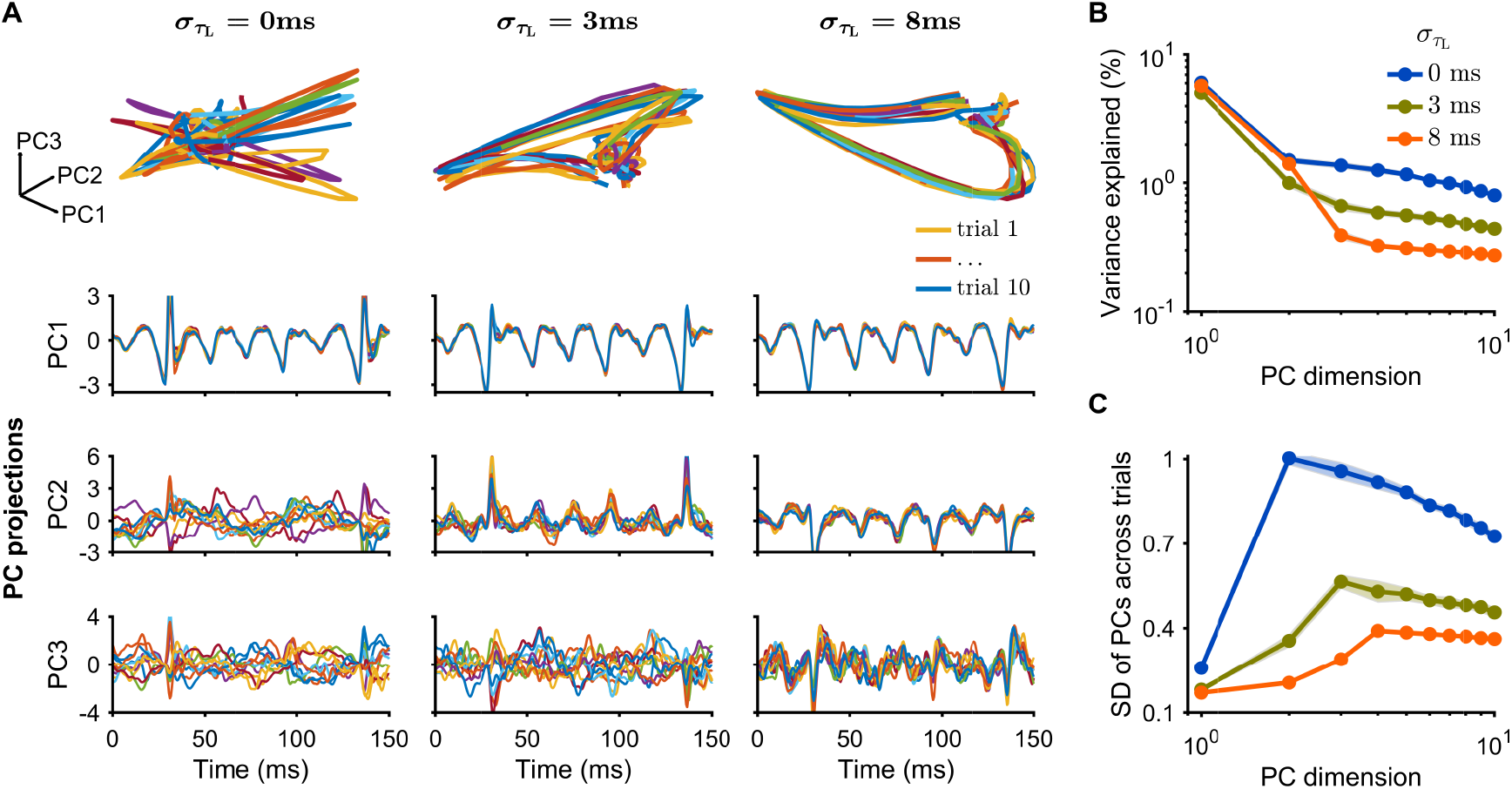
Principal-component analysis of the network activities under MG series input with *f*_MG_ = 50. (**A**) Projections of population responses into three-dimensional space of PC1-PC3 (top) at 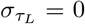 ms (left), 3 ms (middle), 8 ms (right). The overlapped colored lines represent 10 repeated trials. (**B**)Percentages of variance explained by PCs. (**C**) Average trial-to-trial SDs of PCs calculated across 20 trials averaged over time.

Both homogeneous and heterogeneous networks exhibit trial-to-trial stability in the projection onto the first principal component (PC1), reflecting the network-wide response to external input (Fig. 7(A and B)). However, the homogeneous network exhibits considerable trial-to-trial variability in PC2 and PC3 (Fig. 7(A and C)). In contrast, heterogeneous networks (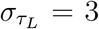 ms and 8 ms) demonstrate stable low-dimensional structures of population responses across trials, indicating a reliable representation of low-dimensional MG input signals (Fig. 7(A and C)). Specifically, the heterogeneous network with higher degree of timescale diversity 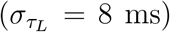 shows markedly reduced trial-to-trial variability in these two components (Fig. 7(A and C)). Thus, heterogeneous networks achieve a stable low-dimensional structure in neural population response activities, and timescale diversity spans larger space with higher dimensions to represent the input in more detail reliably.

The stability of the low-dimensional structure embedded in heterogeneous networks can be attributed to an input-slaved transient dynamics, which enforces the representation robustness. This can be demonstrated by introducing a random perturbation at any time point and examining three subsequent sections of the network’s responses, focusing on three dominant PCs (Fig. 8(A)). During the immediate response to perturbation (section **i**), the low-dimensional structure deviates significantly from that of no perturbation (Fig. 8(A and B)). However, in the subsequent sections **ii** and **iii**, this deviation rapidly converges (Fig. 8(A and B)), indicating the existence of an input-slaved trajectory [27]. The convergence in the low-dimensional structure also informs the performance stability in the MG series reconstruction task (Fig. 8(C)). Thus, networks with input-slaved trajectories can settle into stable, input-induced activity patterns, enhancing the reliability of input signal representation.

**Fig. 8.**
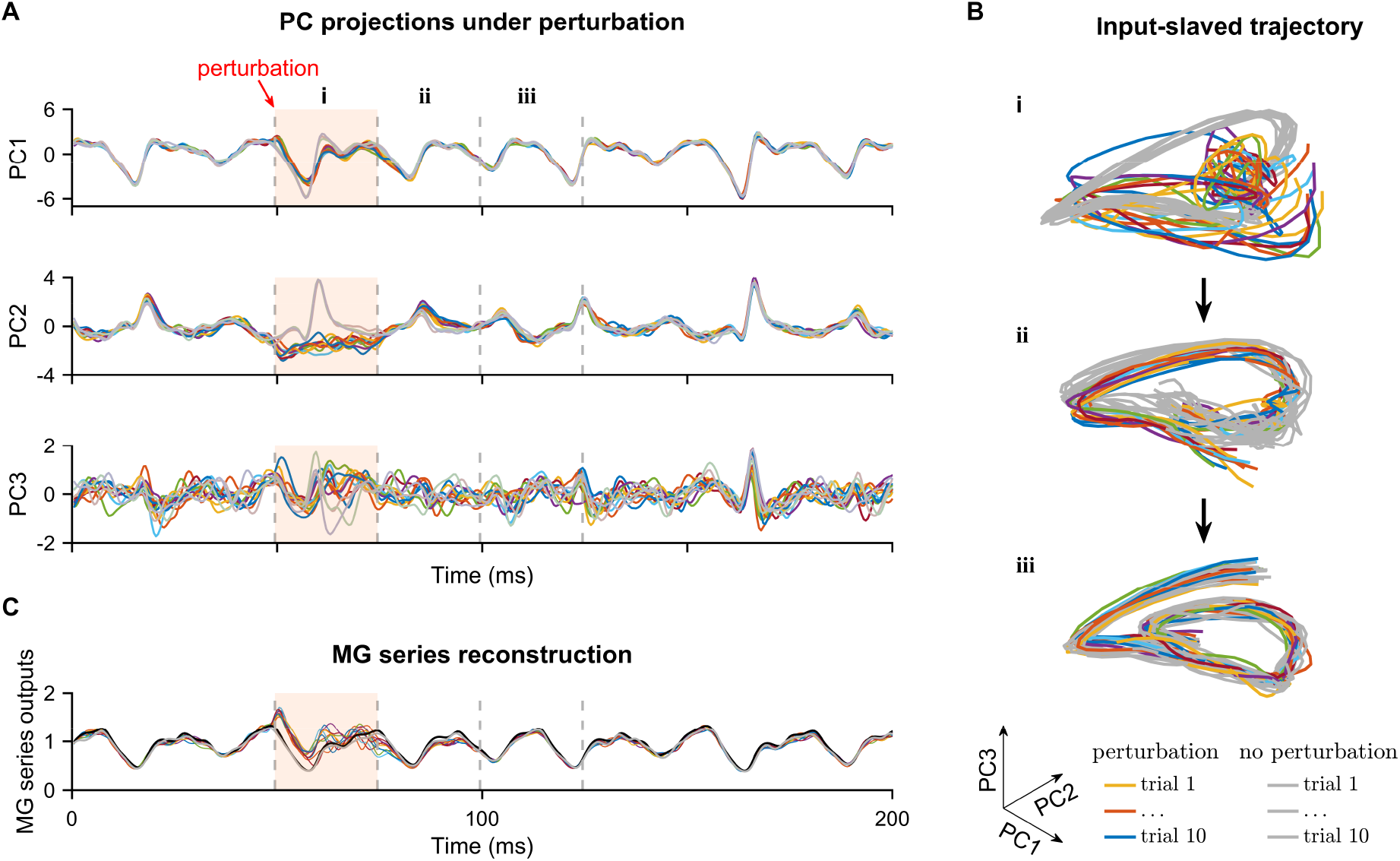
Input-slaved trajectory of neural response under noise perturbation in the heterogeneous network (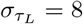 ms). (**A**) Projections of noise-perturbed population responses into PC1, PC2, and PC3. Shaded regions (starting at 50 ms and lasting for 25 ms) indicate the period during which each neuron receives an independent Gaussian noise. (**B**) Projections of perturbed population responses into three-dimensional space, where **i, ii**, and **iii** correspond to three distinct areas delineated by dashed lines in (A), representing the the perturbation period and subsequent intervals. (**C**) MG series reconstructed from network responses to noise perturbation. In (A to C), the overlapping colored lines represent 10 repeated trials with noise perturbation, while grey lines represent trials without perturbation.

These results demonstrate that heterogeneous networks can generate input-slaved trajectories to robustly represent inputs, even though local activity is irregular, variable and sensitive. Therefore, we uncover the underlying dynamical mechanism for achieving reliable neural representation and computation.

### 2.4. This Dynamical Mechanism Explaining Similar Roles of other Neural Heterogeneities

Various neural heterogeneities have significantly contributed to reliable neural computation [13, 16, 17, 22, 23]. We now investigate whether these contributions can be explained by the same dynamical mechanism discussed earlier. As an illustration, we investigate the role of non-uniform input connections, characterized by the correlation range 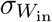 of input connections (details in Sec. 5). It has been shown to enhance representation and computation [13] (also presented in Fig. 9(A)), and it is robust against various connection range ratio *σ*_*i*_*/σ*_*e*_ (data not shown).

**Fig. 9.**
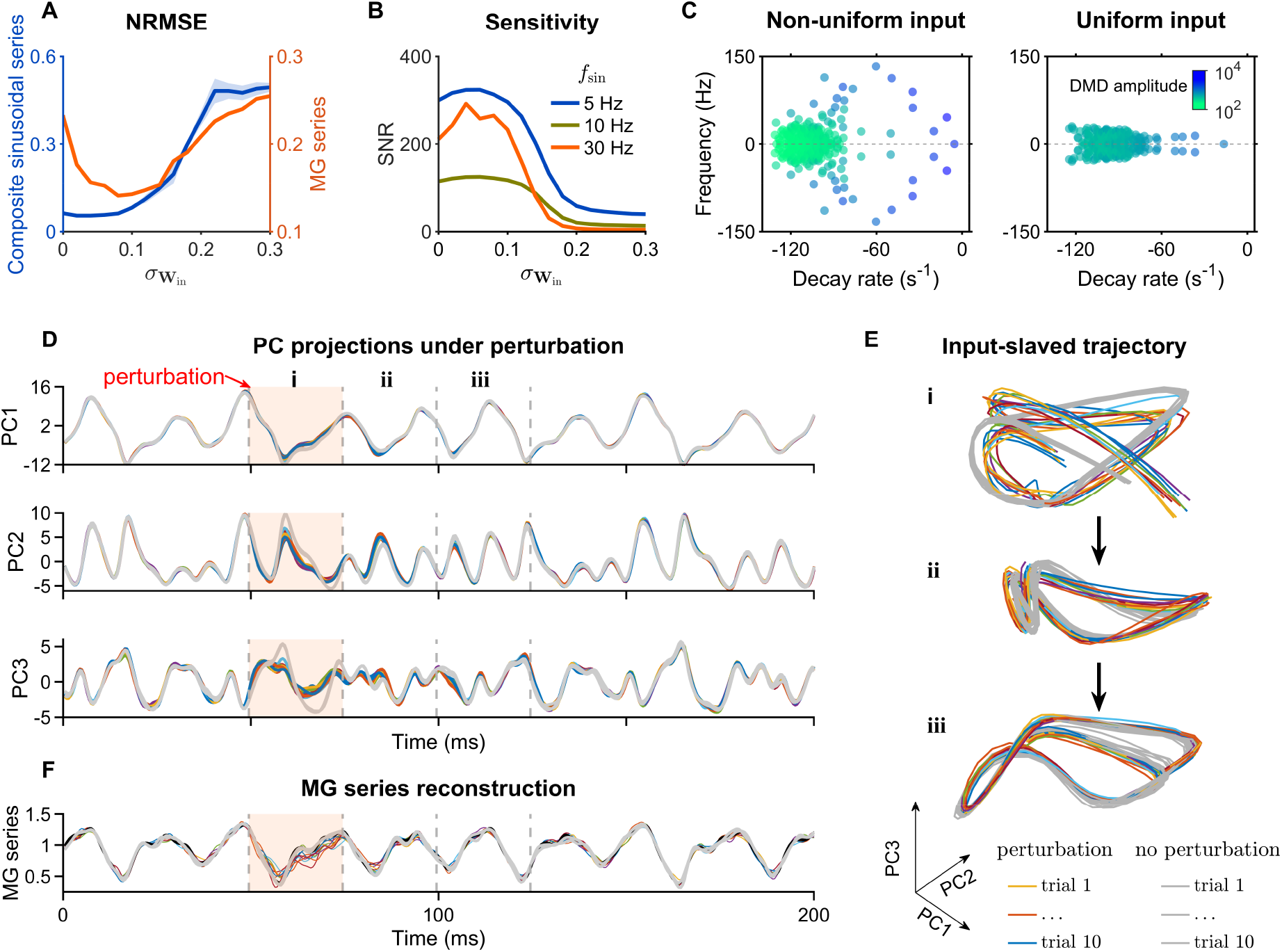
Our mechanism explaining the role of non-uniform input connections in reliable computation. (**A**) NRMSE plotted against 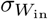 for tasks of input-output mapping (blue line) and MG series reconstruction (orange line). (**B**) Averaged SNR plotted against 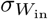 at *f*_sin_ = 5 Hz, 10 Hz, or 30 Hz. (**C**)DMD eigenvalue spectrum of neural responses in networks with non-uniform (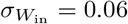, left) or uniform (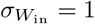, right) input connections. Dot colors represent mode amplitudes. (**D**) Top: Projections of noise-perturbed population responses into PC1, PC2, and PC3. Bottom: MG series reconstructed from network activities under random noise perturbation. (**E**) Perturbed population response in the PC1-PC3 space. In panels (d-f), line colors and shaded regions denote the same as those in Fig. 8. Default time constants are *τ*_*L*_ = 20 ms, *τ*_Γ_ = 20 ms.

Our identified dynamical mechanism can elucidate the role of such input connection heterogeneity: 1) *Disruption of coherent spatiotemporal patterns*: The heterogeneity disrupts the intrinsic coherent spatiotemporal patterns, increasing site activity flexibility and sensitivity (Fig. 9(B)). This allows the network to respond to external stimuli and capture input signal features effectively. 2) *Promotion of consistent stimulus represetation*: It promotes consistent stimulus representation by globally inducing input-related dynamical modes beyond the intrinsic modes (Fig. 9(C)). 3) *Input-slaved trajectories in an expanded representation space*: It expands the representation space to higher dimensions, forming a high-dimensional input-slaved trajectory that reliably represents the input (Fig. 9(D and E)).

As spike threshold heterogeneity also induces firing rate heterogeneity [16, 23] (fig. S7), its effects on network dynamics and function can be accounted for by the same mechanism we have revealed (fig. S8), whereby variations in neuronal properties enhance computation’s reliability. Thus, this mechanism may serve as a general framework for understanding the roles of various neural heterogeneities in reliable representation and computation.

## 3. Discussions

This study elucidates the dynamical mechanism underlying reliable representation and computation in SNNs with neural heterogeneity. In terms of two biologically plausible neuronal timescale diversity, we demonstrate that neural heterogeneity disrupts intrinsic coherent spatiotemporal patterns [13, 26], enhances local sensitivity and aligns neural network activity closely with inputs. This process induces input-related dynamical modes, expands representation space, and forms high-dimensional input-slaved trajectories that reliably represent the input and ensure robust neural information processing against noise, despite neuronal spikings being variable over time and across repeated stimulation trials [35–38]. Thus, we have uncovered a dynamical mechanism that reconciles inherent cortical variability with reliable representation of external inputs. This mechanism also explains the similar roles of other neural heterogeneities, such as non-uniform input connections and spike threshold heterogeneity, serving as a potentially general framework for understanding the role of neural heterogeneity in reliable representation and computation.

### 3.1. Roles of Neural Heterogeneity

Neural heterogeneity, which includes diversity in morphology, type, excitability, connectivity, ion channel distributions and timescales, is ubiquitous in the brain [18–20]. Neuron-to-neuron variability in molecular, genetic, and physiological features is increasingly recognized as a critical aspect of brain function. This biophysical diversity enriches neural system’s dynamical repertoire, fostering individual variability and complex neural dynamics. Complexity and heterogeneity are fundamental to neurobiological systems, manifesting at every level and process, and are intricately linked to the systems’ emergent collective behaviours and functions. Numerous hypothetical roles of neural heterogeneity [39] have been suggested, including: 1) *Population coding*: Enhancing the neuron population’s ability to encode information efficiently [40, 41]; 2) *Reliability*: Increasing the reliability of neural computations [13, 42]; 3) *Working memory*: Supporting complex cognitive functions [43–45] and functional specialization in neural computation [25]; 4) *Reduction in pathological synchronization*: Preventing the onset of pathological states such as epilepsy [23, 46].

Neural heterogeneity provides a richer repertoire of collective dynamics, allowing for more reliable transformations of network inputs [47]. Recent computational studies underscore the importance of spike threshold heterogeneity in enhancing neural network capabilities [16, 25], such as efficient and robust encoding of stimuli [48, 49], improving information transmission and learning [47], and maintaining robustness and persistence of brain function over time and in the face of changing environments (resilience) [23]. Moreover, introducing heterogeneity in the membrane time constants of neurons makes learning complex tasks more stable and robust [17], suggesting that this observed heterogeneity in the brain may be a vital component of the brain’s adaptability.

Despite these findings, previous studies have lacked a dynamical mechanism to understand or unify the roles of various neural heterogeneities, particularly in reconciling inherent cortical variability with reliable representation and computation. Our study uncovers this underlying mechanism beyond current understanding [13, 16, 17, 22–24], opening a new window and providing deeper insights into the role of neural heterogeneity. Our study addresses this gap in understanding by investigating the influence of timescale diversity on network dynamics [50] and its role in shaping reliable neural representation [25]. Our findings highlight the significant role of timescale diversity in cortical functionality and neural computation, focusing on the dissociation of two timescales: leakage time constant *τ*_*L*_ and responsive time constant *τ*_Γ_ [18, 25]. Specifically, our findings show that timescale diversity not only decorates the spatial pattern of collective activities but also broadens the frequency range in neuronal firing patterns [51]. Relaxing one timescale diversity (membrane time constant) to the diversities of two timescales promotes reliable representation and computation, and thus is expected more robust learning. Furthermore, these effects are robust across various connection range ratios, or even randomly shuffled networks.

Our results indicate that neural heterogeneity might contribute to the diversity of receptive fields within a neural population [52], providing a unifying dynamical mechanism for its roles in reliable representation and computation. This mechanism opens new avenues for understanding the complexity and resilience of brain function, suggesting that variations in neuronal properties is crucial for the robustness and adaptability of neural computations.

### 3.2. Input-slaved Transient Dynamics

The brain continuously receives and integrates vast amounts of environmental information, from sensory perception to task-driven behaviors and decision-making processes. Initial studies on neural encoding focused on individual neurons, revealing that certain neurons selectively respond to specific stimulus features such as spatial or temporal frequency, orientation, position, or depth, leading to sparse encoding. However, advancements in recording technologies have shifted research towards the collective dynamics of neural populations [53]. The trajectories of these populations are typically constrained within low-dimensional manifolds in the high-dimensional space of neural activity [9], suggesting that the entire population dynamically encodes stimulus variables in a reduced or coarse-grained neural state space.

Robustness and reproducibility of sequential spatiotemporal responses are essential features of many neural circuits in the sensory and motor systems [9, 54]. Traditional dynamical regimes—such as fixed points, limit cycles, chaotic attractors, and continuous attractors—do not adequately describe reproducible transient sequential neural dynamics. Instead, a stable heteroclinic sequence (SHS) [55, 56], although not an attractor, has been proposed to model these dynamics. SHS trajectories are highly sensitive to input but remain globally stable, with small deviations corrected by distributed stabilizing effects within the neural population. When the input is withdrawn, the system relaxes back to its baseline state. However, SHS is merely an illustrative model, requiring manual design rather than self-organization, and it lacks a solid biophysical basis. Besides, RNNs trained on cognitive tasks have shown that low-dimensional subspaces naturally emerge to support flexible computations at the population level [57, 58]. However, the internal connectivities of these artificial networks do not adhere to biological constraints such as Dale’s principle, which states that neurons must be either excitatory or inhibitory.

Here we discovered a self-organized dynamical mechanism to reliably and robustly produce such input-slaved trajectories for representing the input, based on biologically plausible E-I recurrent SNNs with neural heterogeneity. With no need of pre-designed SHS-like dynamical structures [59], neural heterogeneity disrupts intrinsic coherence between neurons, rendering them sensitive to input for flexible computations. The formed trajectories are transient and nonstationary (Fig. 6(D)), entirely input-induced, and embedded in the complex spatiotemporal neural activities. Despite its distributed coding of the input (fig. S6(C)) and substantial variability of neuronal activities, neural heterogeneity effectively reduces trial-to-trial variability of these trajectories in the noisy system. Thus, ubiquitous neuronal diversity provides a natural biophysical basis for reliable neural information processing.

### 3.3. Impacts on Neuromorphic Computing

Machine learning, particularly through artificial neural networks, has revolutionized computational applications across various fields by excelling in tasks such as regression, classification, prediction, and generation. However, optimizing these networks demands substantial computational resources and energy. Neuromorphic computing [60, 61], which utilizes spiking potentials and waves instead of digital bits, offers a promising alternative by leveraging the inherent efficiencies of biological systems. Recent advancements [62–64] have demonstrated the potential of using nonlinear waves, such as rogue waves and solitons, for neuromorphic computing. These waves’ complex interactions make them particularly suitable for designing reservoir computing. By encoding information onto these waves, it is expected to significantly enhance the development of machine-learning devices leveraging wave dynamics.

Our study adds to this innovative field by elucidating the dynamical mechanism that underpins reliable representation and computation in biologically plausible SNNs characterized by neural heterogeneity. We have discovered that neural heterogeneity disrupts coherent spatiotemporal patterns, thereby enhancing local sensitivity. This mechanism offers the neurons liquid-like superior sensitivity, making them prone to capturing input information efficiently. It also renders the system to generate input-related dynamical modes which are intrinsically nonlinear waves (Fig. 6), thus expressing the complexity necessary for learning from a dataset.

This understanding of neural heterogeneity and its role in reliable computation provides a new avenue for designing new reservoir computing models. By incorporating liquid wave reservoirs, one can leverage the benefits of neuromorphic computing while maintaining the complexity and efficiency needed for high-dimensional data representation and processing. Our discoveries, integrated with the principles of wave dynamics in neuromorphic computing, pave the way for developing advanced reservoir computing that leverages the efficiency and robustness of biological neural information processing.

## 4. Future work

Reliable representation is fundamental to subsequent neural processes such as memory, learning, and decision-making. In this study, basic E-I recurrent SNNs with neural diversity are employed to investigate the underlying dynamical mechanisms of reliable representation and computation. This mechanism, characterized by local sensitivity and global input-slaved transient dynamics due to neural heterogeneity, is expected to persist when the model incorporates more biological details, such as synaptic timescales and plasticity. Specifically, with synaptic timescales mediated by N-methyl-D-aspartate (NMDA) receptor currents, the input-slaved trajectories observed in our study can form the dynamical basis for transient memories, allowing for the temporary storage of information and its flexible use in decision-making [65–67]. Additionally, short-term plasticity can induce traces of hidden states essential for working memory [68–72]. Therefore, our mechanism, which induces high-dimensional input-slaved trajectories, can serve as a substrate for working memory. We plan to investigate these issues further in future.

## 5. Materials and Methods

### 5.1. Network model

In this study, we utilize a spatially extended network of current-based LIF neurons (*N* = 50000), with *N*_*e*_ = 40000 excitatory (**e**) and *N*_*i*_ = 10000 inhibitory (**i**) neurons uniformly spaced on a two-dimensional sheet: Γ = [0, 1] × [0, 1], with torus-like periodic boundaries (Fig. 1(A)). For a neuron located at the coordinate (*i, j*) within population **a** ∈ {**e, i**}, its membrane potential 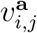 evolves as:

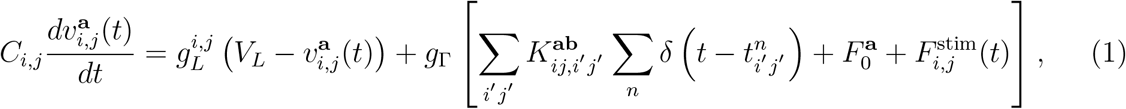

where *V*_*L*_ = −70 mV is the resting potential, *C*_*i,j*_ is the membrane capacitance, 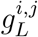 is the leak conductance. The term *g*_Γ_ represents the strength of network drive in the current-based synaptic model, which controls the combined effect of network activity, background input, and external stimulus. 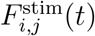 denotes a time-varying external stimulus applied to the neuron, while 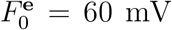 and 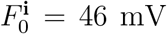 correspond to the constant background input for excitatory and inhibitory neurons, respectively. 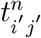 denotes the *n*-th spike time of a neuron located at coordinate (*i*^*′*^, *j*^*′*^) within population **b** ∈ {**e, i**}. When 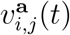 exceeds the spike threshold *V*_th_ = −50 mV, the neuron at coordinate (*i, j*) emits a spike and then its membrane potential is reset to *V*_reset_ = − 75 mV for a refractory period of *τ*_ref_ = 2 ms.

In the mammalian cerebral cortex, neurons establish spatially organized connectivity patterns based on their spatial positions, particularly characterized by widely observed distance-dependent connectivity profiles [12, 73–75]. We utilize a spatially extended network model, as detailed in [13, 28], to approximately model the physiologically observed distance-dependent connectivity among neurons. The synaptic strength from population **b** to population **a** is defined as:

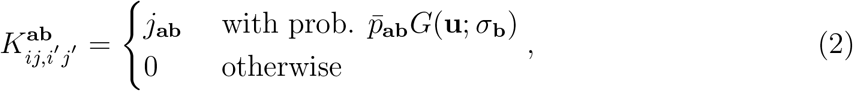

where the two-dimensional wrapped Gaussian function *G*(**u**; *σ*_**b**_) describes the connection probability between neurons at (*i, j*) and (*i*^*′*^, *j*^*′*^). Here, **u** = (*i* − *i*^*′*^, *j* − *j*^*′*^) denotes the spatial distance between the neurons, and the standard deviation of Gaussian function *σ*_**b**_ determines the spatial width of excitatory and inhibitory synaptic projections. Unless stated otherwise, we use the excitatory and inhibitory projection width: *σ*_**e**_ = 0.05 and *σ*_**i**_ = 0.1. The average connection probabilities for excitatory and inhibitory populations are 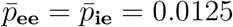 and 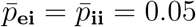 with synaptic weights of *j*_**ee**_ = 2 mV, *j*_**ie**_ = 4 mV, and *j*_**ei**_ = *j*_**ii**_ = −5 mV.

### 5.2. Timescale diversity and spike threshold diversity

Cortical neurons exhibit remarkable diversity in both morphological features and electrophysiological properties. Data-driven modeling based on the Allen Cell Types Database reveals a wide distribution of parameters, including capacitance and conductance, among cortical neurons [76]. In this study, we considered two distinct types of time constants contributing to timescale diversity within the LIF neuron model [18, 25] (Fig. 1(B)).

Leakage time constant, denoted as 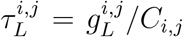, characterizes the neuronal timescale for membrane potential evolving to resting potential in the absence of network input. On the other hand, response time constant, denoted as 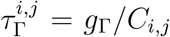, represents the timescale of the neuronal response to incoming stimuli. Leakage timescale diversity arises from diverse membrane capacitance and conductance among neurons, while response timescale diversity reflects diverse membrane capacitance. To model the experimentally observed long-tail distributions of neuronal parameters determining neuronal timescales [18], we employed a log-normal distribution of neuronal timescales with its probability density function *ξ*_*m*_ expressed as:

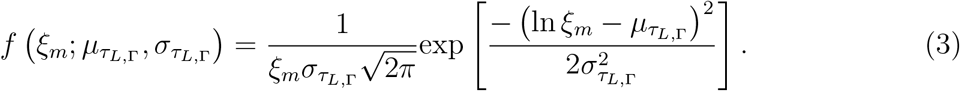

Here, *ξ*_*m*_ (*m* = *L*, Γ, *C*) are independent from each other, 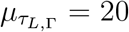 ms represent the mean of the *τ*_*L*_ (or *τ*_Γ_) distribution, and 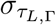 denotes the parameter controlling the degree of variability in the *τ*_*L*_ (or *τ*_Γ_) distribution. Higher values of 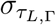 indicate a greater degree of timescale diversity.

According to the defined relationship, distributions of leakage timescale *τ*_*L*_ and response timescale *τ*_Γ_ can be correlated, denoted as *ρ*(*τ*_*L*_, *τ*_Γ_). When *τ*_*L*_ and *τ*_Γ_ are fully correlated (*τ*_*L*_ = *τ*_Γ_ and *ρ* = 1), the model is equivalent to that used in the previous study, which employed a single parameter *τ*_*m*_ to control neuronal timescale [17]. In this study, we sampled *τ*_*L*_ and *τ*_Γ_ from a common set to generate correlated distributions. Specifically, we randomly drew a proportion of 1 − *ρ* from the distribution *ξ*_*L*_ (or *ξ*_Γ_), while the remaining proportion *ρ* was drawn from the distribution *ξ*_*C*_. Thus, when *ρ* = 0, *τ*_*L*_ and *τ*_Γ_ are independent, whereas *ρ* = 1 indicates complete correlation between *τ*_*L*_ and *τ*_Γ_.

In addition, we also employed a lognormal distribution for spike threshold diversity when considering other types of neural heterogeneities:

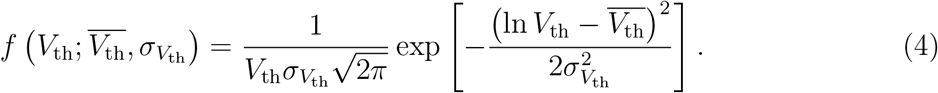

Here, the mean 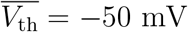, and the standard deviation 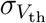 controls the degree of spike threshold diversity.

### 5.3. External Inputs

In this study, we examined the external time-varying input received by each neuron within the network. This input is defined as 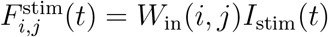, where *W*_in_(*i, j*) denotes the input weight of a neuron located at coordinate (*i, j*), and *I*_stim_(*t*) represents the external input signal. We considered two types of input weight distributions: (1) uniform input connections, where *W*_in_(*i, j*) is constant at 1, and (2) non-uniform input connections, where the weight varies spatially. For the latter, we generated *W*_in_ via a spatially correlated Gaussian noise with a mean of zero and a covariance function expressed as:

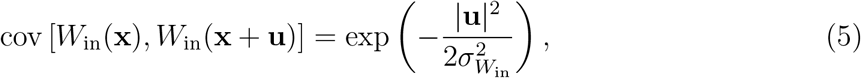

where |**u**| = |(*i* − *i*^*′*^, *j* − *j*^*′*^)| denotes the spatial distance. The parameter 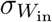 determines the spatial correlation range of *W*_in_ in the two-dimensional space. The weight distribution of *W*_in_ becomes increasingly spatially heterogeneous as 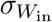 decreases.

Two types of time series were utilized as external driving signals: a sinusoidal periodic series and a Mackey-Glass chaotic series. The sinusoidal periodic series is mathematically defined as:

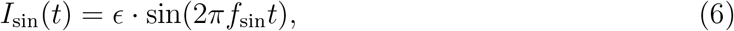

where *f*_sin_ and *ϵ* represent the frequency and amplitude of the sinusoidal stimulus signal, respectively. The Mackey-Glass (MG) chaotic time series is represented as:

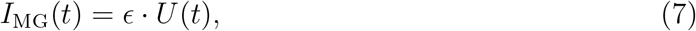

with the amplitude *ϵ* set to 20 mV. The variable *U* (*t*) is governed by the Mackey–Glass equation:

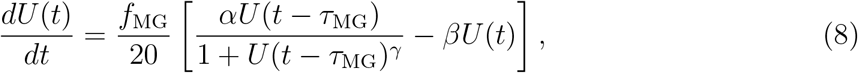

where the parameters *α* = 0.2, *β* = 0.1, and *γ* = 10 are dimensionless and define the system dynamics along with a time delay of *τ*_MG_ = 17 ms. The parameter *f*_MG_ is pivotal in controlling the pace of the Mackey–Glass dynamics. A higher value of *f*_MG_ accelerates the evolution of the Mackey–Glass equation within a specified duration, thereby regulating the frequency of the quasi-periodic Mackey–Glass series.

### 5.4. Response robustness under noise perturbation

To examine the noise robustness of the network’s response to external MG series input, we additionally applied a period of independent Gaussian noise to each neuron:

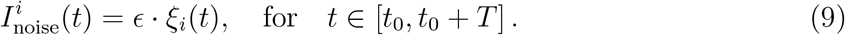

Here, *ξ*_*i*_(*t*) ∼ 𝒩 (0, 1) with covariance cov [*ξ*_*i*_(*t*), *ξ*_*j*_(*t*)] = *δ*_*ij*_. The duration *T* = 25 ms and the amplitude *ϵ* = 40 mV.

### 5.5. Measurements

#### 5.5.1. Instantaneous neural firing rate

In our study, we recorded coarse-grained neural firing rates *R*(*t*) from 2500 sites across the whole network, with each recording site comprising *N*_local_ = 16 excitatory neurons. For each recording site, the site firing rate *R*(*t*) was calculated as follows:

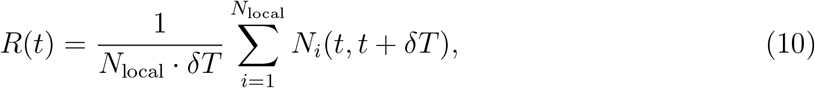

where *N*_*i*_(*t, t* + *δT*) represents the spike count of neuron *i* within a time bin of *δT* = 0.5 ms.

#### 5.5.2. Synchrony index

To quantify the collective spiking synchronization of neurons within each recording site, we employed the synchrony index based on the averaged Pearson correlation coefficient of spike counts between neuron pairs. The synchrony index is defined as follows:

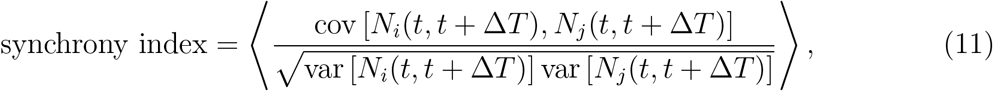

where *N*_*i*_(*t, t* + Δ*T*) and *N*_*j*_(*t, t* + Δ*T*) denote the spike counts of neurons *i* and *j* within a time window of Δ*T* = 100 ms. Covariances (cov) and variances (var) of these spike counts were computed across the windows. The notation ⟨·⟩ denotes averaging across all neuron pairs, excluding the cases where *i* = *j*.

#### 5.5.3. Spectrum analysis and signal-to-noise ratio

The power spectrum *S*(*f*) of site firing rates was calculated using the fast Fourier transform (FFT), and the mean power spectrum was then derived by averaging the power spectra across all recording sites: ⟨*S*(*f*)⟩. The time-frequency spectrum of the MG series was calculated using the complex Morlet wavelet transform, with a bandwidth of 3.0 and a wavelet center frequency of 0.5 Hz.

The signal-to-noise ratio (SNR) was used to quantify the strength of the network’s response to external periodic stimuli. Based on the power spectral density *S*(*f*) of neural response activity, the SNR was calculated using the following equation:

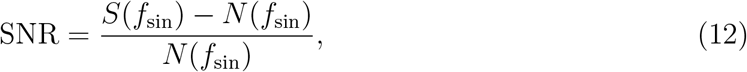

where *S*(*f*_sin_) denotes the power at the frequency *f*_sin_, and *N* (*f*_sin_) is the averaged power at nearby frequencies within a small range around *f*_sin_ (∼ 1 Hz). A high SNR value indicates a strong response of neural activity to the external periodic stimulus at the frequency *f*_sin_.

#### 5.5.4. Least-square method

In our study, we extracted coarse-grained neural firing rates *X* (with a recording site number *n* = 2500) from network response activities and used them to reconstruct the output signal **ŷ** = *W*_out_*X*. The readout weight matrix *W*_out_ that minimizes the difference between the reconstructed output **ŷ** least-square method (LSM):

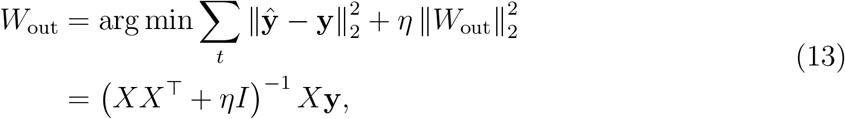

and the target (desired) signal **y** is obtained through the ridge where ∥·∥_2_ is the Frobenius norm, *I* is the matrix, and *η* = 100 is the regularization parameter that penalizes large values in the parameter vector *W*_out_. We explored the range of *η* from 0 to 200, finding that it does not significantly impact the main results of our study.

#### 5.5.5. Dynamic Mode Decomposition

We utilized the Dynamic Mode Decomposition (DMD) algorithm, a data-driven eigen-decomposition method, to evaluate the sensitivity of network response to Mackey-Glass (MG) signal input. DMD directly approximates the observed system dynamics from high-dimensional data within a specified time interval, enabling the extraction of a low-dimensional representation and decomposing complex dynamic systems into coherent spatiotemporal modes [34, 77]. The DMD procedure constructs a locally linear dynamical system proxy that transfers the network’s current state *X* to its subsequent state *X*^*′*^:

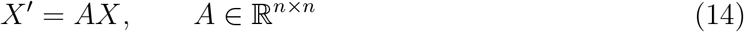

where

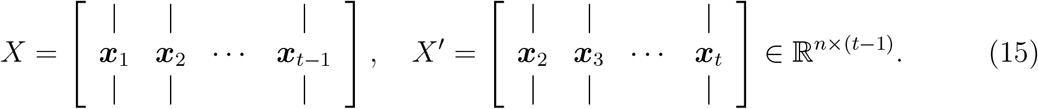

Here, ***x***_*t*_ represents the vector of coarse-grained neural firing rates (*n* = 2500) at time point *t*. The optimal linear operator *A* is determined by minimizing the Frobenius norm error ∥ *X*^*′*^− *AX*∥_*F*_. In practical application, the DMD algorithm involves the following key steps [77]:

1. Decompose neural population firing data, denoted as *X*, using Singular Value Decomposition (SVD) with rank-*r* truncation: 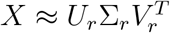.
2. Calculate the matrix representation Ã by projecting the full matrix *A* onto the Proper Orthogonal Decomposition (POD) space: 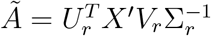.
3. For comparison with the power spectrum obtained through FFT, scale the DMD mode amplitudes as 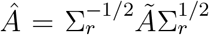, and perform eigendecomposition: Ã = *Q*Λ*Q*^*T*^, where *Q* and Λ are the eigenvector and diagonalizable eigenvalue matrices, respectively.
4. Calculate DMD modes as 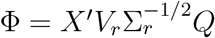.
5. Approximate the system’s future state as 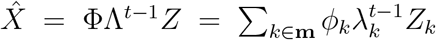, where *Z* = Φ^†^ *X*_0_. Here, the notation † denotes the pseudo-inverse of the Φ matrix, the set **m** = {1, 2, · · ·, *r*} represents the labels of the selected DMD modes, and *X*_0_ represents the initial state of the system.

In this study, we specifically selected DMD modes to predict the system’s short-term future state at 4 time steps ahead (*t* = 4 ms). Subsequently, using the least squares method, we reconstructed the Mackey-Glass (MG) output signal based on the predicted future state 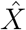. The performance of the reconstructed signal serves as an indicator of which DMD modes effectively capture the dynamic features of the MG input signal. In essence, the presence of DMD modes capable of accurately predicting the input MG signal suggests a high responsivity of the network to the input signal.

#### 5.5.6. Reliability of neural representation

The reliability of the neural representation under external stimuli was quantified using the normalized root-mean-square error (NRMSE) in the test dataset:

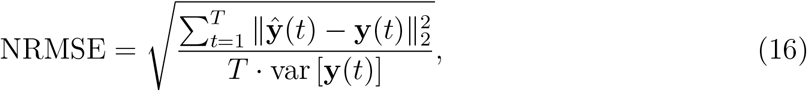

where *T* is the number of time steps and var [**y**(*t*)] denotes the variance of the target signal. A low NRMSE value indicates a robust alignment between the reconstructed signal and the target sequence across multiple trials, reflecting consistent neural representations achieved by the network. Thus, a lower NRMSE value indicates higher reliability in the network representation. Here, the training and testing datasets consist of 50 and 10 trials of neural response data, respectively, with each trial lasting for a 500 ms stimulation duration.

#### 5.5.7. Largest Lyapunov exponent

Lyapunov exponents measure the sensitivity of a nonlinear system’s dynamics to initial conditions, with positive Lyapunov exponents indicating chaotic behavior. Here, we compute the largest Lyapunov exponent (LLE) based on site firing rate series to characterize the flexibility of neural activity. A smaller Lyapunov exponent suggests that neural dynamics are closer to a steady state or periodic orbit, whereas a larger Lyapunov exponent indicates greater divergence from the initial dynamical trajectory over time. Thus, a larger Lyapunov exponent corresponds to more flexibility and variability in spontaneous neural activity over time. To estimate the LLE from highly variable firing rate series, we used the state space reconstruction with principal components (SSRPC) algorithm and the Python-based SSRPC toolbox proposed by [78], which improves noise robustness compared to the original Rosenstein’s algorithm [79].

### 5.6. Computer simulations

For numerical integration of the equations, we employed the Euler method, except for the Mackey-Glass equation, for which we used the 4th-order Runge-Kutta method. The simulation time step was set to *dt* = 0.05 ms. To ensure statistically reliable results, we performed 20 independent simulations.

## Author contributions

D.Y. and S.W. designed the study; S.W. performed the experiments and analyses; All authors contributed to writing the manuscript.

## Data and materials availability

All data are available in the main text or the supplementary materials. Computer code for all simulations and analysis of the resulting data is available from the corresponding author upon reasonable request.

## 6. Supplementary Materials

**Fig. S1.**
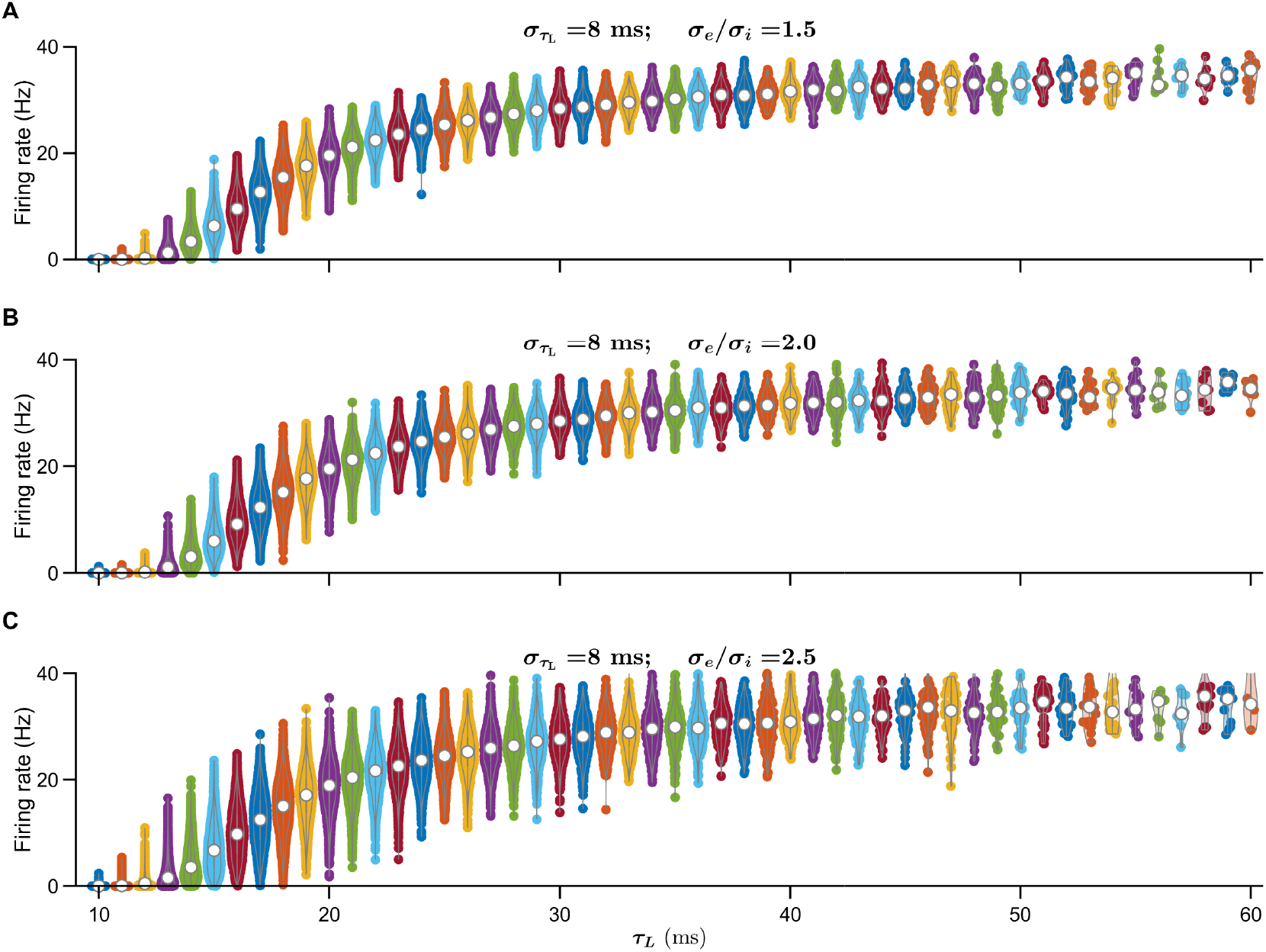
Timescale-dependent Gaussian distributions of firing rate. Violin plots of firing rate distributions for recording sites with identical timescales *τ*_*L*_ in the heterogeneous network 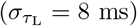 at various ratios: *σ*_*e*_*/σ*_*i*_ = 1.5 (A), 2.0 (B), or 2.5 (C).

**Fig. S2.**
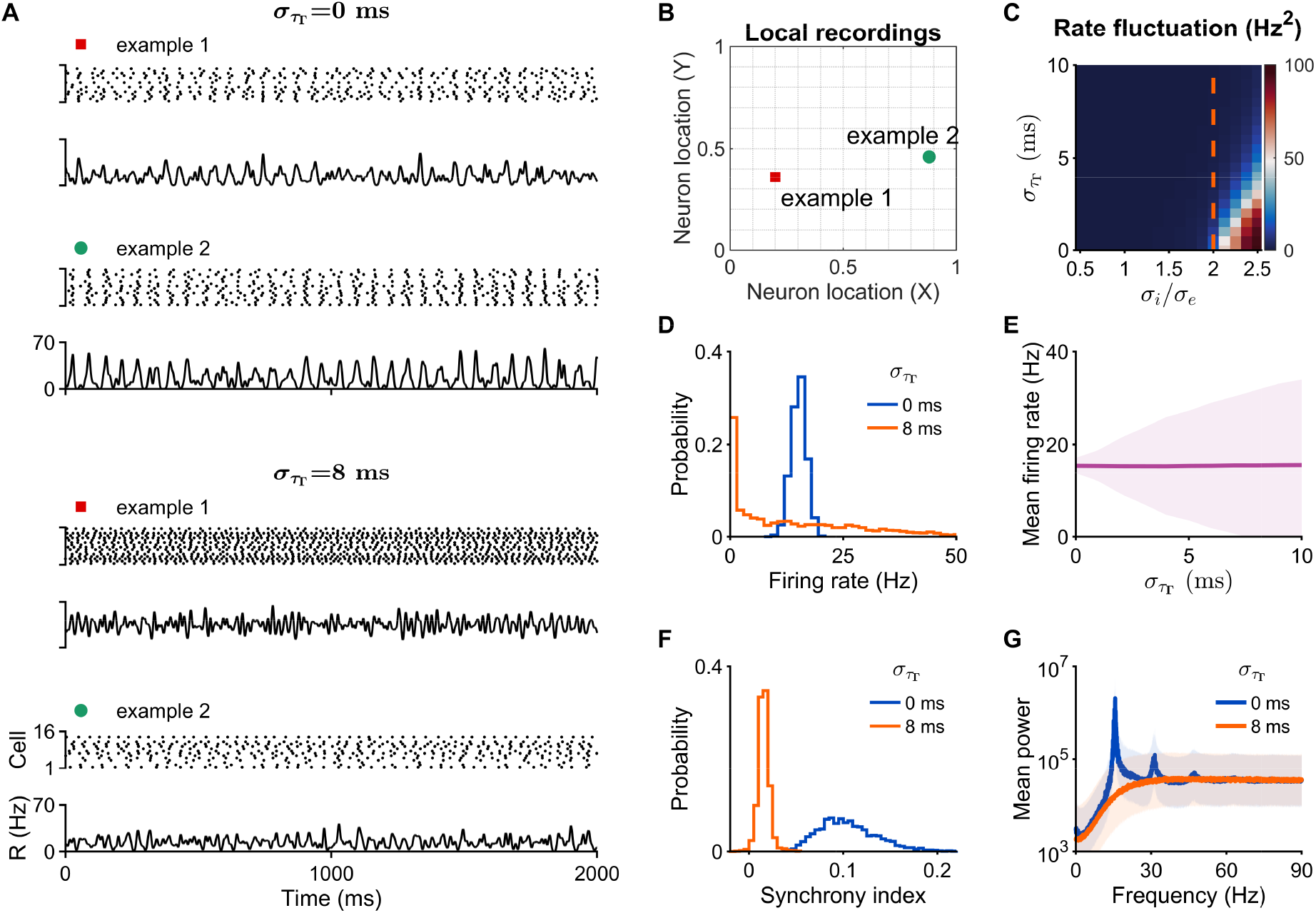
Spontaneous spatiotemporal dynamics in spatially extended networks with/without *τ*_Γ_ diversity. (**A**) Raster plot and firing rate series of two randomly chosen spatial sites in the homogeneous network (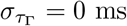, top) or the heterogeneous network (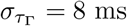, bottom). (**B**) Spatial positions of the two randomly chosen sites within the network. (**C**) Temporal fluctuation of E neurons’ firing rates across the parameter space (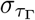, *σ*_*i*_*/σ*_*e*_). (**D**) Firing rate distributions of E neurons at 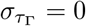 or 8 ms. (**E**) Mean firing rates and their SDs (shaded areas) plotted against 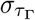. (**F**) Synchrony index distributions. (**G**) Average power spectrum of individuals’ firing rates at 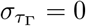 ms or 8 ms.

**Fig. S3.**
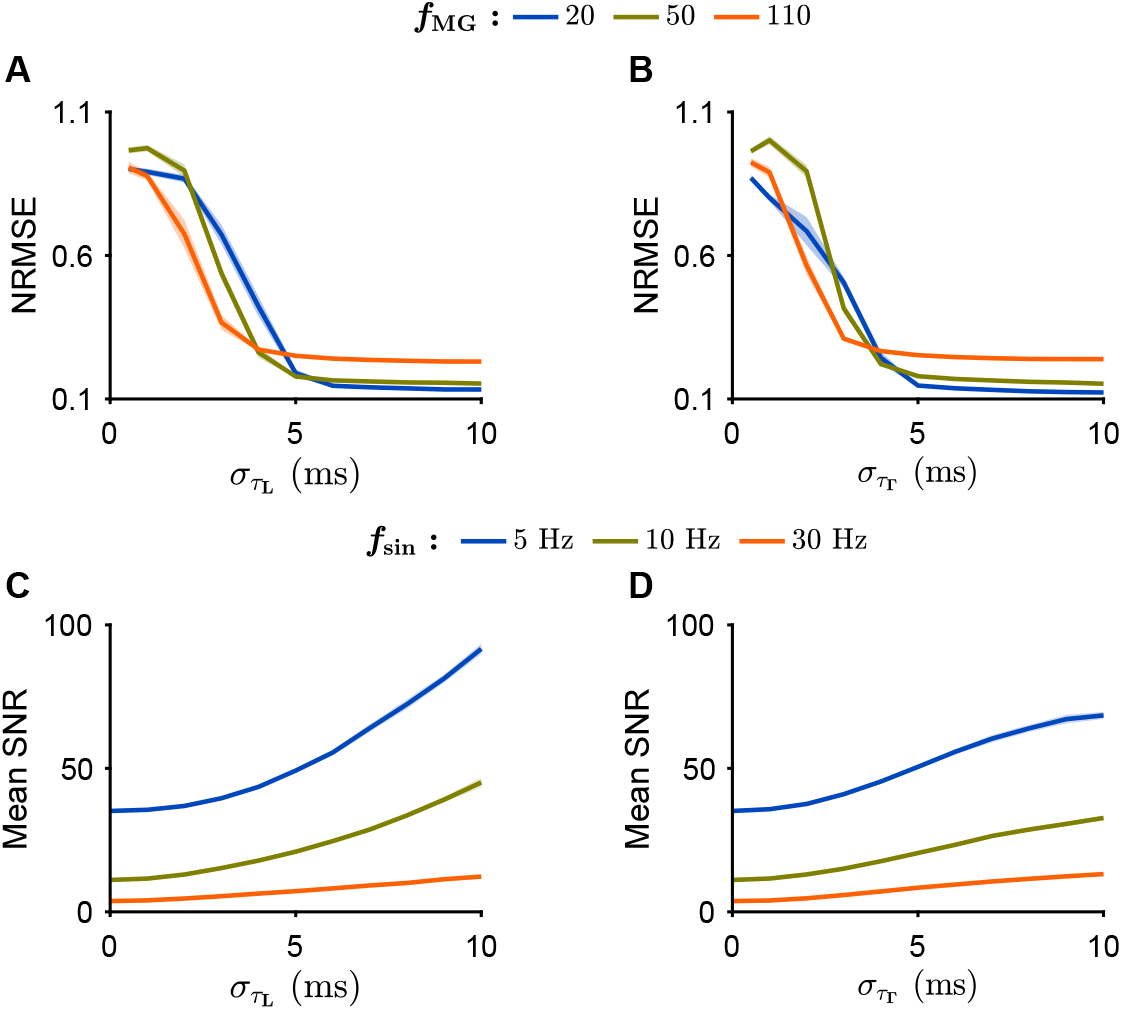
Results extended to networks with randomly shuffled connections. (**A** and **B**) NRMSE plotted against 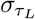 (A) and 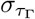 (B) at *f*_MG_ = 20, 50, or 110. (**C** and **D**) Mean SNR plotted against 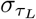 (C) and 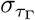 (D) at *f*_sin_ = 5 Hz, 10 Hz, or 30 Hz.

**Fig. S4.**
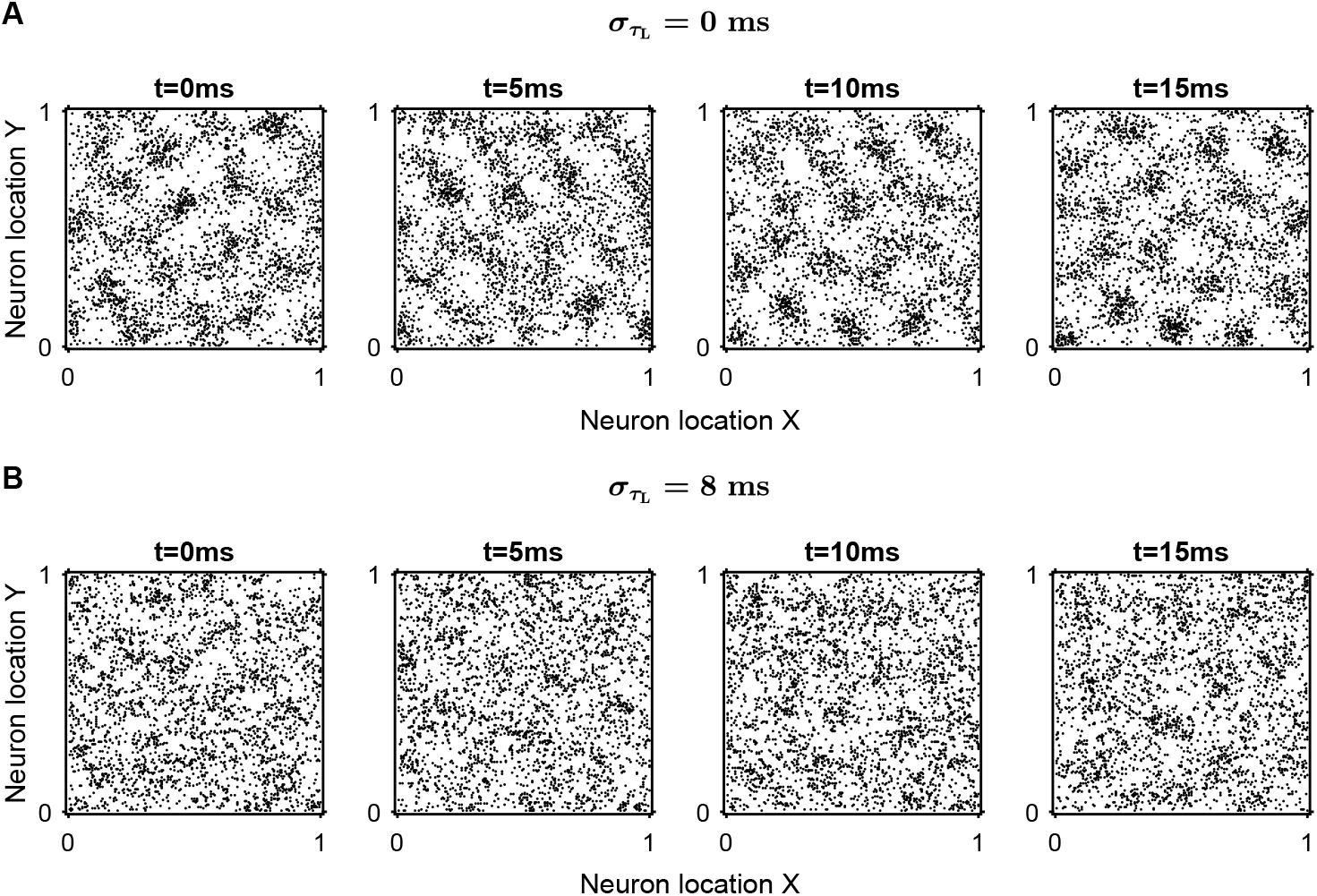
Comparison of excitatory firing patterns between networks with and without timescale diversity. (**A** and **B**) Snapshots of spiking activity in spatial networks with 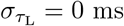 (A) and 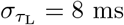 (B) over 5 ms time windows at time points: *t* = 0 ms, 5 ms, 10 ms, and 15 ms.

**Fig. S5.**
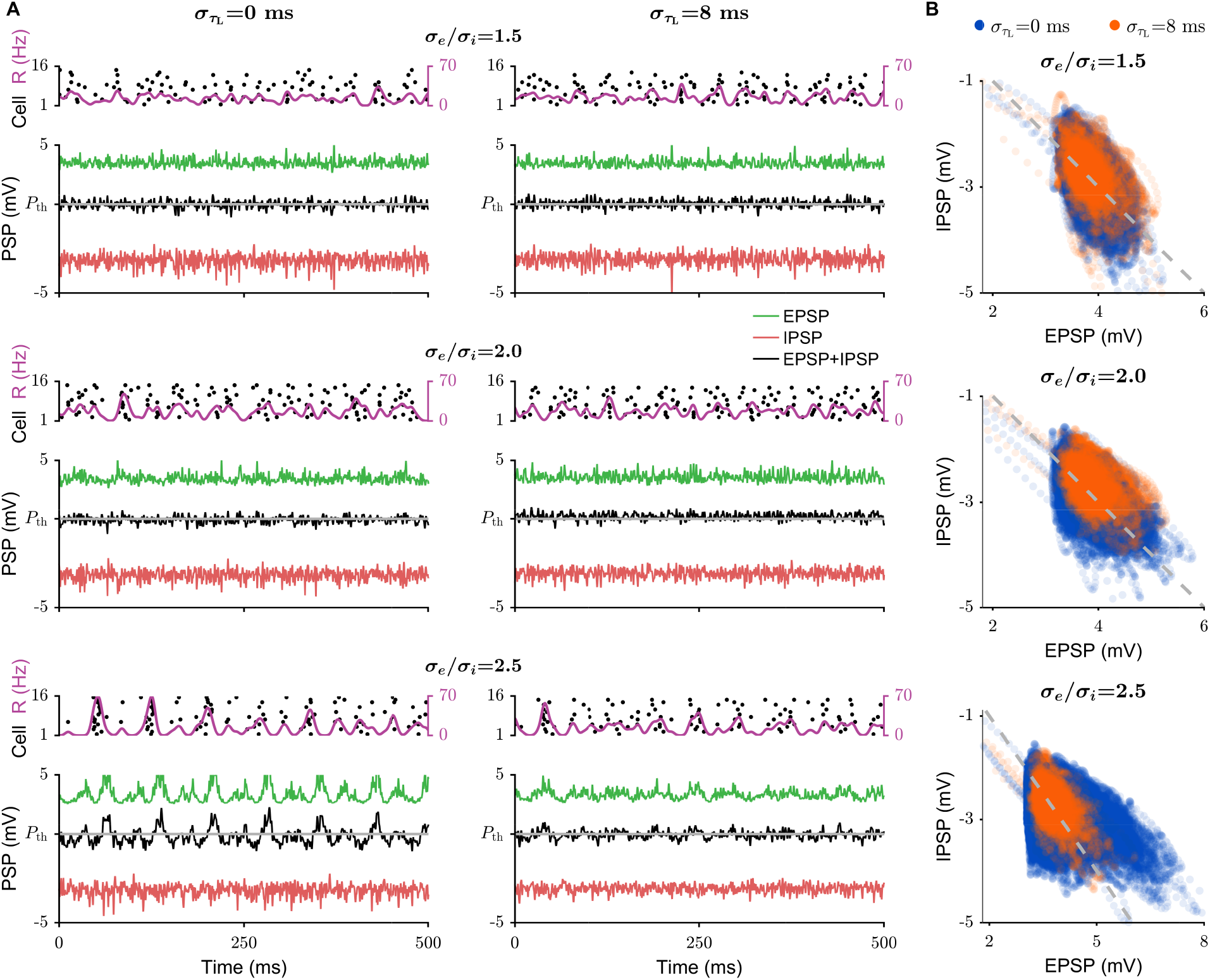
Timescale diversity regulates E-I balance in spatially extended networks. (**A**) Firing activity and averaged postsynaptic potentials (PSP) recorded from sites in homogeneous 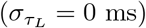 and heterogeneous 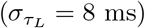 networks at various ratios: *σ*_*i*_*/σ*_*e*_ =1.5, 2.0, and 2.5. The purple lines represent the traces of firing rates, while the green, red, and black lines correspond to the E postsynaptic potential (EPSP), I postsynaptic potential (IPSP), and their combination (EPSP + IPSP), respectively. (**B**) Scatter plots of EPSP against IPSP for homogeneous and heterogeneous networks with different *σ*_*i*_*/σ*_*e*_. In panels(A) and (B), the grey lines represent the postsynaptic potential threshold *P*_th_ above which neurons generate action potentials (spikes).

**Fig. S6.**
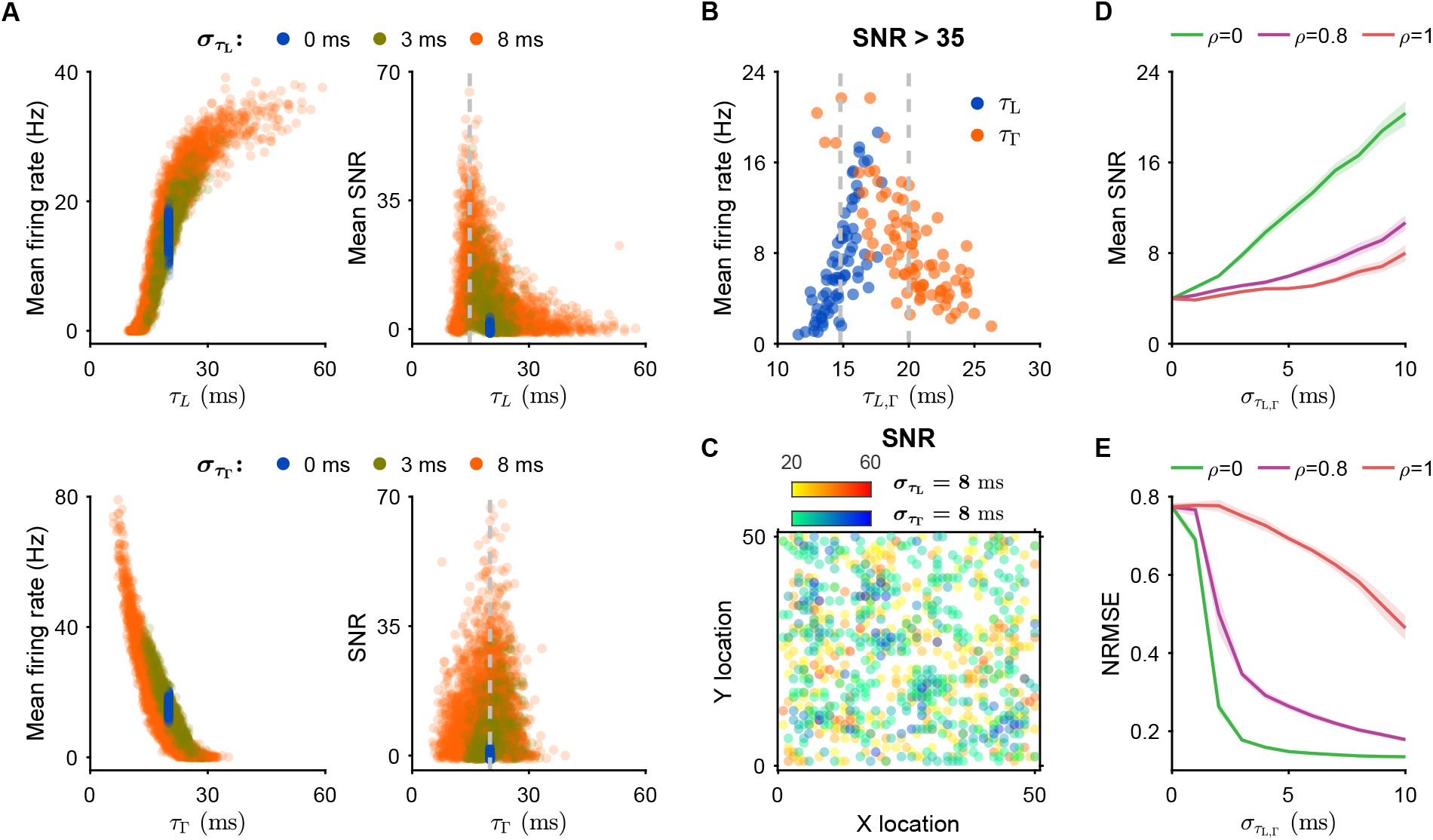
Correlation between leakage and response timescales impedes the beneficial effects of timescale diversity on computation’s reliability. (**A**) Mean firing rates of spontaneous network activities (left) and SNRs of network responses (right) plotted against *τ*_*L*_ (top) or *τ*_Γ_ (bottom) for networks with (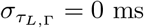, 3 ms, or 8 ms). (**B**) Mean firing rates of network responses plotted against *τ*_*L*_ (blue) or *τ*_Γ_ (orange) for recording sites with SNRs larger than 35. (**C**) Spatially distributed neural response (SNR *>* 20). Red and blue dots represent the recording sites in networks with 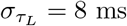 and 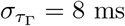, respectively. Deeper colors indicate higher SNR values. (**D** and **E**) Mean SNR (D) and NRMSE (E) plotted against 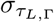 with various correlations of *τ*_*L*_ and *τ*_Γ_: *ρ* = 0.0, 0.8, and 1.0. In panels (A) and (B), the dashed lines represent the mean of *τ*_*L*_ or *τ*_Γ_ for recording sites with SNRs larger than 35. The frequency and amplitude of the sinusoidal input are respectively *f*_sin_ = 30 Hz and *ϵ* = 3 mV in panels (a-d). In panel (E), *f*_MG_ = 50.

**Fig. S7.**
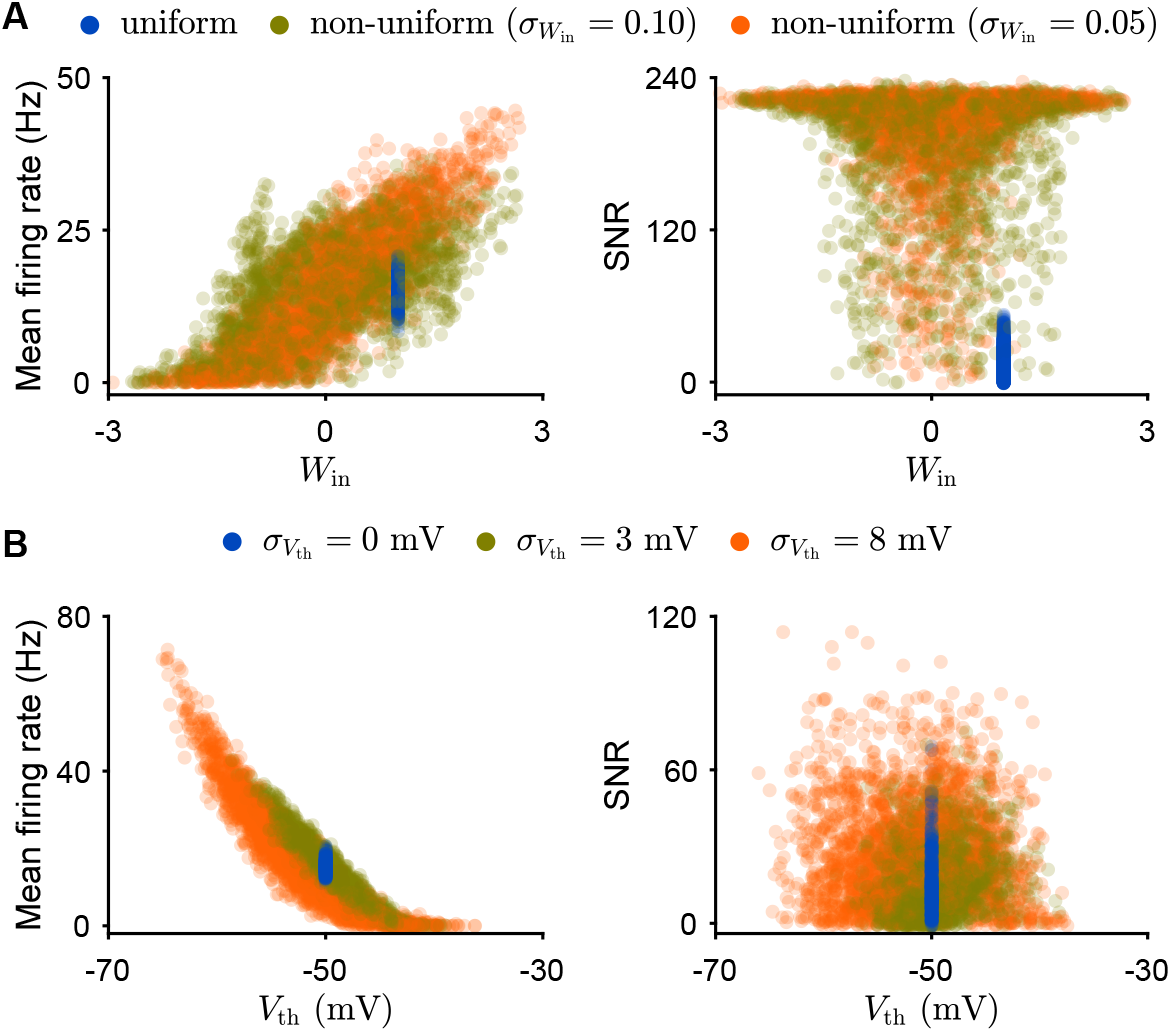
Response heterogeneity in networks with other types of diversity. (**A** and **B**) Mean firing rates of spontaneous network activities (left) and SNR of network responses (right) plotted against *W*_*in*_ for various degrees of non-uniform input connections (A) and against spike thresholds *V*_th_ for different levels of spike threshold diversity (B, 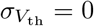 mV, 3 mV, and 8 mV). In both panels, networks receive a constant external input in the spontaneous state and a sinusoidal input in the response state. In panel (b, left), the input amplitude is *ϵ* = 0 mV; in all other cases, *ϵ* = 3 mV. The sinusoidal input frequency is *f*_sin_ = 10 Hz.

**Fig. S8.**
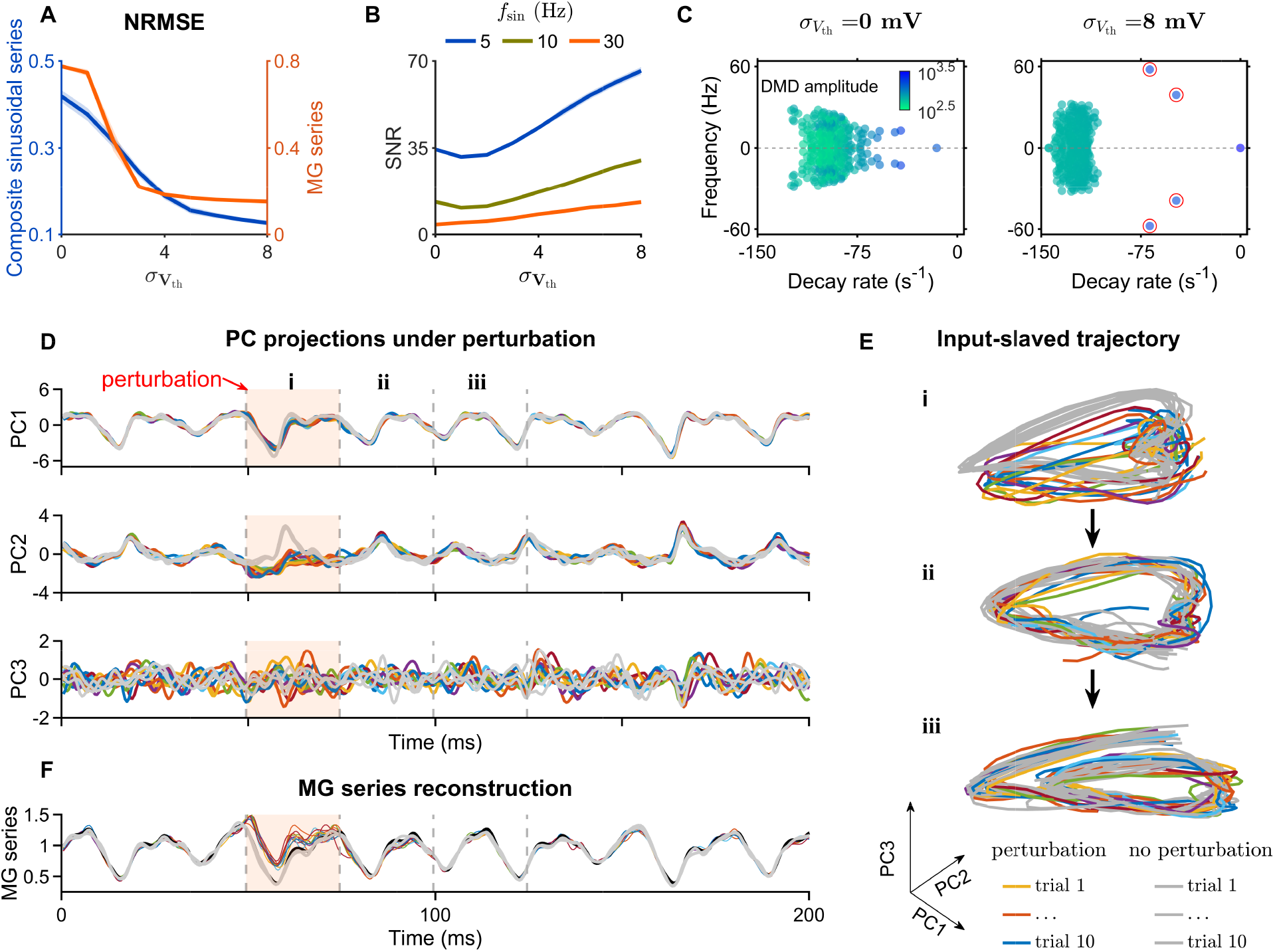
Our mechanism explaining the role of spike threshold heterogeneity in reliable computation. (**A**) NRMSE plotted against *V*_th_ for tasks of input-output mapping (blue line) and MG series reconstruction (orange line). (**B**) Averaged SNR plotted against *V*_th_ at *f*_sin_ = 5 Hz, 10 Hz, and 30 Hz. (**C**) DMD eigenvalue spectrum for neural responses in networks with 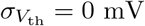 (left) and 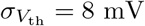 (right) under MG series input. Dot colors represent mode amplitudes and red circled blue dots represent the input-induced two new dominant modes. (**D** to **F**) Population responses to noise perturbation in the network with spike threshold heterogeneity. (D) Projections of noise-perturbed population responses into PC1, PC2, and PC3. (E) Perturbed population responses in the PC1-PC3 space. (F) MG series reconstructed from the network under random noise perturbation. The overlapping colored lines represent 10 repeated trials with noise perturbation, while the grey lines represent trials without perturbation. Shaded regions (starting at 50 ms and lasting for 25 ms) indicate periods during which independent Gaussian noise perturbations were applied to each neuron. Default time constants are *τ*_*L*_ = 20 ms and *τ*_Γ_ = 20 ms.

